# Dopamine neuron morphology and output are differentially controlled by mTORC1 and mTORC2

**DOI:** 10.1101/2021.11.07.467637

**Authors:** Polina Kosillo, Kamran M. Ahmed, Bradley M. Roberts, Stephanie J. Cragg, Helen S. Bateup

## Abstract

The mTOR pathway is an essential regulator of cell growth and metabolism. Midbrain dopamine neurons are particularly sensitive to mTOR signaling status as activation or inhibition of mTOR alters their morphology and physiology. mTOR exists in two distinct multiprotein complexes termed mTORC1 and mTORC2. How each of these complexes affect dopamine neuron properties and whether they act together or independently is unknown. Here we investigated this in mice with dopamine neuron-specific deletion of *Rptor* or *Rictor*, which encode obligatory components of mTORC1 or mTORC2, respectively. We find that inhibition of mTORC1 strongly and broadly impacts dopamine neuron structure and function causing somatodendritic and axonal hypotrophy, increased intrinsic excitability, decreased dopamine production, and impaired dopamine release. In contrast, inhibition of mTORC2 has more subtle effects, with selective alterations to the output of ventral tegmental area dopamine neurons. As mTOR is involved in several brain disorders caused by dopaminergic dysregulation including Parkinson’s disease and addiction, our results have implications for understanding the pathophysiology and potential therapeutic strategies for these diseases.

## Introduction

The mechanistic target of rapamycin (mTOR) is an evolutionarily conserved kinase that serves as a central coordinator of cellular metabolism and regulator of anabolic and catabolic processes (Saxton and Sabatini, 2017). Balanced mTOR signaling is required for proper cell growth and function, while dysregulation of mTOR signaling is associated with various diseases (Crino, 2016; Karalis and Bateup, 2021; Saxton and Sabatini, 2017). Within the nervous system, mTOR fulfills distinct functions at different developmental stages. During embryonic development, mTOR regulates progenitor cell proliferation, differentiation, and neuronal migration (Blair and Bateup, 2020; Magri and Galli, 2013; Switon et al., 2017). In neurons, mTOR controls morphology, physiology, and synaptic properties (Costa-Mattioli and Monteggia, 2013; Hoeffer and Klann, 2010; Switon et al., 2017). Consequently, dysregulation of mTOR signaling has profound impact on nervous system function and several neurodevelopmental, psychiatric and neurodegenerative disorders are directly caused by or associated with altered mTOR activity (Costa-Mattioli and Monteggia, 2013; Karalis and Bateup, 2021; Lipton and Sahin, 2014).

While mTOR is present and active in all cells, distinct neuronal types show differential responses to alterations in mTOR signaling (Benthall et al., 2021; Yang et al., 2012). Midbrain dopamine (DA) neurons residing in the substantia nigra pars compacta (SNc) and ventral tegmental area (VTA) are particularly sensitive to mTOR signaling status, which may be a result of their unique physiology and high metabolic demands (Matsuda et al., 2009; Pacelli et al., 2015). For example, adult deletion of *mTOR* from VTA DA neurons leads to decreased DA release in the nucleus accumbens (NAc), altered DA transporter (DAT) expression, and attenuated synaptic and behavioral responses to cocaine (Liu et al., 2018b). Treatment with the mTOR inhibitor rapamycin, applied directly to striatal slices, decreases DA axon terminal size and depresses evoked DA release (Hernandez et al., 2012). By contrast, chronic activation of mTOR signaling via deletion of genes encoding the upstream negative regulators Pten or Tsc1 leads to DA neuron hypertrophy and increased DA synthesis, with differential effects on DA release (Diaz-Ruiz et al., 2009; Kosillo et al., 2019). In Parkinson’s disease (PD) models, both partial mTOR inhibition and mTOR activation can be neuroprotective to degenerating DA neurons via different mechanisms. These include activation of autophagy, suppression of pro-apoptotic protein synthesis, enhancement of neuronal survival, and increased axon growth (Zhu et al., 2019). Therefore, up or down-regulation of mTOR signaling can impact multiple aspects of DA neuron biology. Since perturbations in dopaminergic mTOR signaling are implicated in neurodevelopmental and psychiatric disorders (Dadalko et al., 2015b; Kosillo et al., 2019), degeneration and neuroprotection in PD (Cheng et al., 2011; Dijkstra et al., 2015; Kim et al., 2012; Malagelada et al., 2010; Ries et al., 2006) and drug-induced neuroplasticity and tolerance (Collo et al., 2013; Liu et al., 2018b; Mazei-Robison et al., 2011), it is important to understand how mTOR controls DA neuron cytoarchitecture and function.

mTOR participates in two multi-protein complexes termed mTORC1 and mTORC2, which have both shared and unique components (Liu and Sabatini, 2020; Switon et al., 2017). These complexes have distinct upstream activators and downstream targets, although some signaling crosstalk has been observed (Liu and Sabatini, 2020; Xie and Proud, 2014). Manipulations used to study mTOR signaling are often not specific to one complex. Rapamycin treatment can interfere with both mTORC1 and mTORC2 signaling (Karalis, 2021; Sarbassov et al., 2006), deletion of *Pten* activates both mTORC1 and mTORC2 (Chen et al., 2019), and loss of Tsc1 activates mTORC1 but suppresses mTORC2 signaling (Karalis, 2021). Consequently, from the current literature it is difficult to disentangle which mTOR complex is responsible for the observed effects. For mTOR-related diseases, it will be important to define which mTOR complex is most relevant for pathophysiology and which should be targeted in a potential therapeutic approach.

The functions of mTORC1 have been well-studied and in many cell types it controls protein synthesis by regulating the phosphorylation of downstream targets including 4E-BP1 and p70S6K kinase (Burnett et al., 1998; Fingar, 2002; Gingras et al., 2001). P70S6K in turn phosphorylates ribosomal protein S6, a canonical readout of mTORC1 activity. In neurons, alterations in mTORC1 signaling result in changes in neuronal morphology, cell metabolism, intrinsic excitability, synaptic transmission and plasticity (Costa-Mattioli and Monteggia, 2013; Switon et al., 2017). The functions of mTORC2 are less well understood; however, emerging evidence indicates that it controls somatodendritic architecture, synaptic properties and certain forms of synaptic plasticity (Angliker et al., 2015; Angliker and Ruegg, 2013; McCabe et al., 2020; Urbanska et al., 2012; Zhu et al., 2018). Ser473 on Akt is the most well-studied target of mTORC2 signaling, which also phosphorylates and controls the activity of PKC and GSK3 (Baffi et al., 2021; Sarbassov et al., 2005). Notably, manipulations that alter mTORC1 or mTORC2 signaling can lead to both overlapping and distinct phenotypes in neurons and these can vary by cell type (Angliker et al., 2015; McCabe et al., 2020; Urbanska et al., 2012). Therefore, a careful dissection of the relative roles of mTORC1 and mTORC2 in different types of neurons is important for understanding how this central signaling pathway controls neuronal structure and function, in both health and disease.

Our group has previously shown that constitutive activation of mTORC1 in DA neurons, due to deletion of the gene encoding the upstream negative regulator Tsc1, leads to pronounced somatodendritic hypertrophy, reduced intrinsic excitability, and strongly impaired DA release despite increased DA synthesis (Kosillo et al., 2019). Remarkably, we found that complete suppression of mTORC1 signaling via deletion of the gene encoding the obligatory mTORC1 component Raptor (Hara et al., 2002; Kim et al., 2002), was as detrimental to DA neurotransmission as loss of Tsc1. However, partially constraining mTORC1 activity with heterozygous loss of *Rptor* was sufficient to prevent impaired DA release in the context of Tsc1 loss. Thus, balanced mTORC1 signaling is necessary for proper dopaminergic output while aberrant mTORC1 activity is detrimental. However, the mechanism by which suppression of mTORC1 caused DA release deficits in Tsc1 KO DA neurons is unknown.

While mTORC1 is a known regulator of cell size (Fingar, 2002), including in DA neurons (Kosillo et al., 2019), it was shown that mTORC2 and not mTORC1 is responsible for somatic hypotrophy in VTA DA neurons in response to chronic morphine (Mazei-Robison et al., 2011). In this study, the morphine effects on VTA DA neuron soma size and intrinsic excitability could be phenocopied by deletion of *Rictor*, an obligatory binding partner of mTORC2 (Sarbassov et al., 2004), and prevented by *Rictor* overexpression (Mazei-Robison et al., 2011). Other studies showed alterations in DA tissue content and DAT function in mice with *Rictor* deleted from excitatory or catecholaminergic neurons (Dadalko et al., 2015b; Siuta et al., 2010). Therefore, open questions remain regarding the specific DA neuron properties that are regulated by mTORC1 and mTORC2 and whether these complexes act together or independently.

Here we addressed this by directly comparing how DA neuron-specific deletion of *Rptor* or *Rictor* affects key cellular properties of SNc and VTA DA neurons. We find that disruption of mTORC1 strongly impacted DA neuron structure and function leading to global cellular hypotrophy, increased intrinsic excitability, reduced DA synthesis, and impaired DA release. By contrast, suppression of mTORC2 generally resulted in more mild morphological changes and selectively increased the excitability but reduced the output of VTA DA neurons. Disruption of both mTOR complexes by concomitant deletion of *Rptor* and *Rictor* led to pronounced deficits that were greater than with suppression of either complex alone, suggesting that phenotypes resulting from inhibition of mTORC1 and mTORC2 may arise via independent mechanisms.

## Results

### Somatodendritic architecture of DA neurons is altered by mTORC1 or mTORC2 inhibition

The activity of mTORC1 or mTORC2 can be suppressed by deletion of the genes encoding their respective obligatory components Raptor or Rictor (Figure 1a). To achieve DA neuron-specific inhibition of each mTOR complex, we crossed DAT-IRES-Cre (Bäckman et al., 2006) (DAT-Cre) mice to *Rptor^fl/fl^* (Peterson et al., 2011; Sengupta et al., 2010) or *Rictor^fl/fl^* (Magee et al., 2012) conditional knock-out (KO) mice (Supplemental Figure S1a-b). In this model, Cre expression turns on around embryonic day 17.5 (Bäckman et al., 2006). Deletion of *Rictor* or *Rptor* at this time point did not impact DA neuron differentiation or survival as the total numbers of tyrosine hydroxylase (TH)-expressing DA neurons in the midbrain at 8-12 weeks of age were not different between *Rptor^fl/fl^;DAT-Cre^wt/+^* (DA-Raptor KO) or *Rictor^fl/fl^;DAT-Cre^wt/+^* (DA-Rictor KO) mice and their respective wild-type (WT) littermate controls (Supplemental Figure S1c-h).

**Figure 1.**
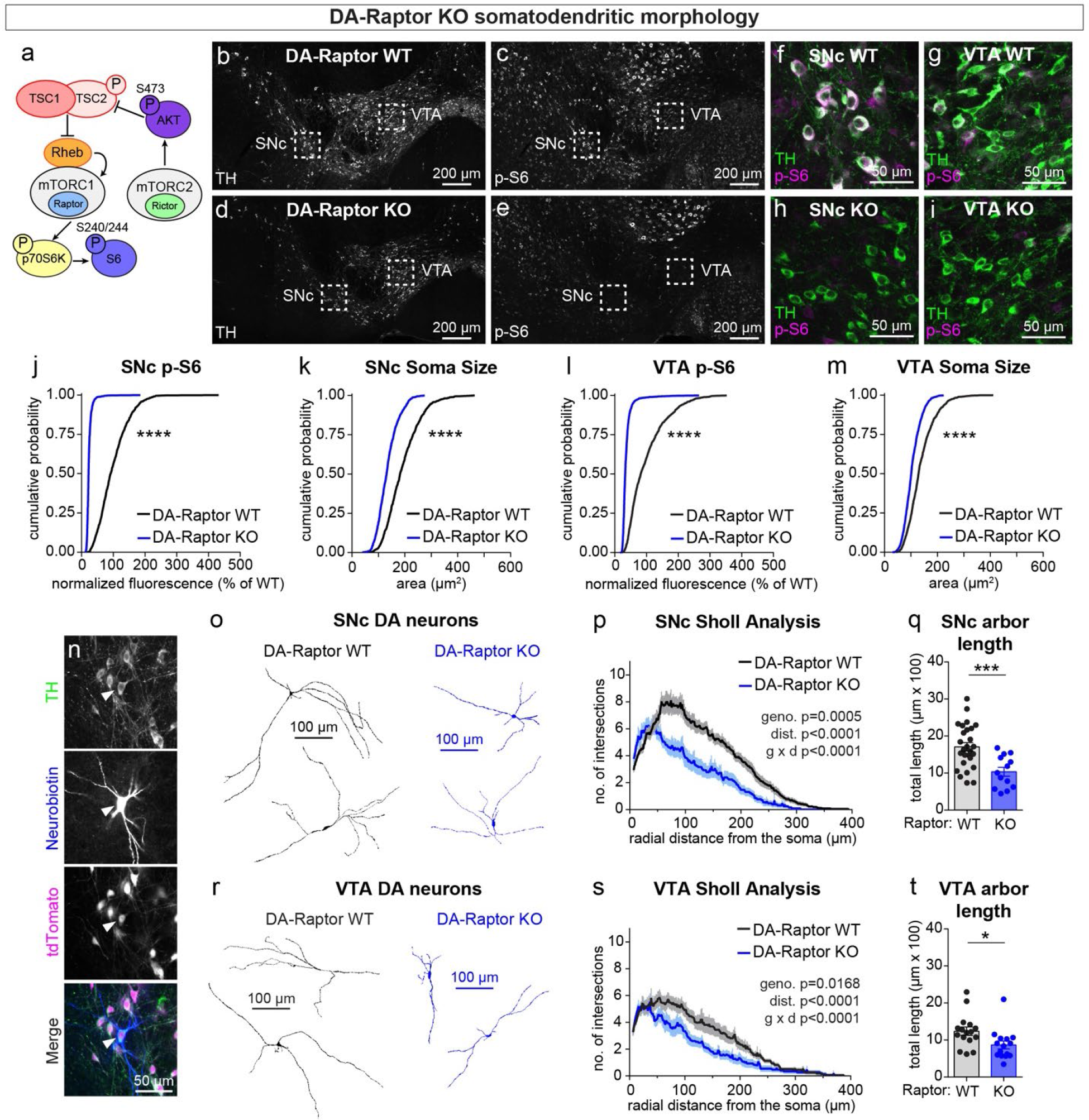
mTORC1 suppression causes somatodendritic hypotrophy of DA neurons. **a)** Simplified mTOR signaling schematic showing mTORC1 and mTORC2 with their obligatory components Raptor and Rictor, respectively. Ribosomal protein S6 is phosphorylated on Ser240/244 by p70S6K, a direct phosphorylation target of mTORC1. AKT is phosphorylated on Ser473 by mTORC2. AKT is an upstream regulator of the Tsc1/2 complex, which negatively regulates mTORC1 activity via the small GTPase Rheb. **b-e)** Representative confocal images of midbrain sections from DA-Raptor WT (**b.c**) and DA-Raptor KO (**d,e**) mice labeled with antibodies against tyrosine hydroxylase (TH) and phosphorylated S6 (p-S6, Ser240/244); scale bars=200 μm. **f-i)** Higher magnification merged images of the SNc (**f,h**) and VTA (**g,i**) boxed regions from panels in b-e; scale bars=50 μm. **j,k)** Cumulative distributions of SNc DA neuron p-S6 levels (**j**) and soma area (**k**). DA-Raptor WT in black: n=1024 neurons from three mice, DA-Raptor KO in blue: n=1045 neurons from three mice; ****p<0.0001, Kolmogorov–Smirnov tests. **l,m)** Cumulative distributions of VTA DA neuron p-S6 levels (**l**) and soma area (**m**). DA-Raptor WT in black: n=1389 neurons from three mice, DA-Raptor KO in blue: n=1526 neurons from three mice; ****p<0.0001, Kolmogorov–Smirnov tests. **n)** Representative confocal images of a midbrain section containing a triple-labelled DA neuron (TH, neurobiotin and tdTomato Cre-reporter; arrowhead) used for dendritic arbor reconstruction and analysis. **o)** Reconstructions of the dendrites and cell body of SNc DA neurons of the indicated genotypes. **p)** Sholl analysis of SNc DA neuron dendritic arbors. Dark colored lines are the mean, lighter color shading is the SEM. DA-Raptor WT in black: n=27 neurons from 9 mice, DA-Raptor KO in blue: n=13 neurons from 6 mice. Two-way ANOVA p values are shown. **q)** Mean ± SEM total dendritic arbor length per cell. DA-Raptor WT: n=27 neurons from 9 mice, DA-Raptor KO: n=13 neurons from 6 mice; ***p=0.0003, Welch’s two-tailed t-test. Dots represent values from individual neurons. **r)** Reconstructions of the dendrites and cell body of VTA DA neurons of the indicated genotypes. **s)** Sholl analysis of VTA DA neuron dendritic arbors. Dark colored lines are the mean, lighter color shading is the SEM. DA-Raptor WT in black: n = 16 neurons from 6 mice, DA-Raptor KO in blue: n=15 neurons from 6 mice. Two-way ANOVA p values are shown. **t)** Mean ± SEM total dendritic arbor length per cell. DA-Raptor WT: n = 16 neurons from 6 mice, DA-Raptor KO: n = 15 neurons from 6 mice; *p=0.0235, Welch’s two-tailed t-test. Dots represent values from individual neurons. See also (Supplemental Figure 1).

SNc and VTA DA neurons are distinct in terms of their synaptic inputs and projection targets (Beier et al., 2015; Lammel et al., 2011; Watabe-Uchida et al., 2012), morphological and electrophysiological properties (Lammel et al., 2008), and vulnerability to disease (Brichta and Greengard, 2014; Gantz et al., 2018). We therefore examined the impact of mTOR signaling perturbations on SNc and VTA DA neuron populations separately. We observed the expected inhibition of mTORC1 signaling in DA-Raptor KO mice, shown by strongly reduced phosphorylation of S6 (p-S6) and decreased soma size of both SNc and VTA DA neurons (Figure 1b-m). Since reductions in Raptor can alter dendritic morphology in other neuron types (Angliker et al., 2015; Urbanska et al., 2012), we filled individual midbrain DA neurons with neurobiotin (Figure 1n) and performed 3D reconstructions to assess their dendritic morphology. Consistent with findings in other neuron types, DA-Raptor KO neurons in the SNc and VTA exhibited significantly reduced total dendrite length and dendritic complexity as determined by Sholl analysis, with more pronounced changes in SNc neurons (Figure 1o-t). Thus, constitutive mTORC1 suppression leads to significant hypotrophy of midbrain DA neurons characterized by reduced soma size and dendritic arborization.

We have previously shown that Cre-dependent deletion of *Rictor* reduces p-473 Akt levels in cultured hippocampal neurons, indicative of suppressed mTORC2 activity (Karalis, 2021). Due to technical limitations of the antibody, we were not able to assess p-473 at the level of individual neurons in brain sections. To assess mTORC1 signaling status in DA-Rictor KO mice, we measured p-S6 levels and found a small but significant reduction in p-S6 in both SNc and VTA DA neurons compared to WT controls (Figure 2a-i,k). Although p70S6K and S6 are not direct phosphorylation targets of mTORC2 (see Figure 1a), this observation is consistent with previous reports that mTORC2 suppression can lead to reduced activity of mTORC1 (McCabe et al., 2020; Urbanska et al., 2012), suggesting some crosstalk between mTORC1 and mTORC2. Consistent with their reduced S6 phosphorylation, DA-Rictor KO neurons in the SNc and VTA exhibited a small but significant reduction in soma size (Figure 2j,l). However, unlike mTORC1 suppression, inhibition of mTORC2 increased the number of proximal dendritic branches measured by Sholl analysis, while the total dendrite length of SNc and VTA DA neurons was unchanged (Figure 2m-r). Together, these data show that mTORC2 suppression reduces mTORC1 activity in DA neurons, which is associated with a small but significant decrease in soma size. In contrast to mTORC1 inhibition, suppression of mTORC2 increases proximal dendrite branching.

**Figure. 2.**
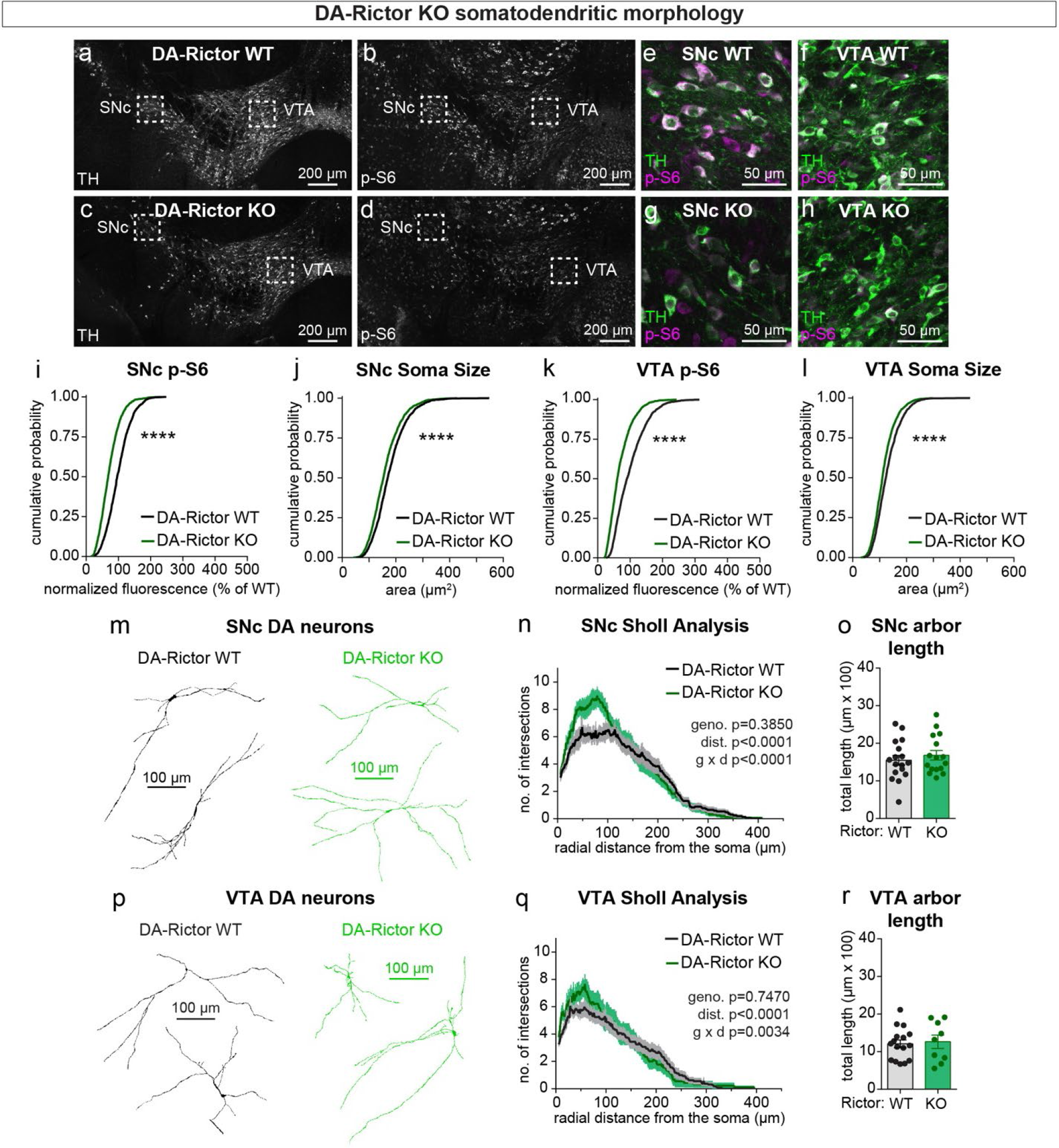
mTORC2 inhibition reduces DA neuron soma size and increases proximal dendrite branching. **a-d)** Representative confocal images of midbrain sections from DA-Rictor WT (**a,b**) and DA-Rictor KO (**c,d**) mice labeled with antibodies against tyrosine hydroxylase (TH) and phosphorylated S6 (p-S6, Ser240/244); scale bars=200 μm. **e-h)** Higher magnification merged images of the SNc (**e,g**) and VTA (**h,i**) boxed regions from panels in b-e; scale bars=50 μm. **i,j)** Cumulative distributions of SNc DA neuron p-S6 levels (**i**) and soma area (**j**). DA-Rictor WT in black: n=1280 neurons from three mice, DA-Rictor KO in green: n=1550 neurons from four mice; ****p<0.0001, Kolmogorov–Smirnov tests. **k,l)** Cumulative distributions of VTA DA neuron p-S6 levels (**k**) and soma area (**l**). DA-Rictor WT in black: n=1968 neurons from three mice, DA-Rictor KO in green: n=2370 neurons from four mice; ****p<0.0001, Kolmogorov–Smirnov tests. **m)** Reconstructions of the dendrites and cell body of SNc DA neurons of the indicated genotypes. **n)** Sholl analysis of SNc DA neuron dendritic arbors. Dark colored lines are the mean, lighter color shading is the SEM. DA-Rictor WT in black: n=17 neurons from 6 mice, DA-Rictor KO in green: n=16 neurons from 6 mice. Two-way ANOVA p values are shown. **o)** Mean ± SEM total dendritic arbor length per cell. DA-Rictor WT: n=17 neurons from 6 mice, DA-Rictor KO: n=16 neurons from 6 mice; p=0.4633, Welch’s two-tailed t-test. Dots represent values from individual neurons. **p)** Reconstructions of the dendrites and cell body of VTA DA neurons of the indicated genotypes. **q)** Sholl analysis of VTA DA neuron dendritic arbors. Dark colored lines are the mean, lighter color shading is the SEM. DA-Rictor WT in black: n=16 neurons from 7 mice, DA-Rictor KO in green: n=9 neurons from 5 mice. Two-way ANOVA p values are shown. **r)** Mean ± SEM total dendritic arbor length per cell. DA-Rictor WT: n = 16 neurons from 7 mice, DA-Rictor KO: n = 9 neurons from 5 mice; p=0.7907, Welch’s two-tailed t-test. Dots represent values from individual neurons. See also (Supplemental Figure 1).

### Striatal dopamine axon density is reduced by mTORC1 or mTORC2 inhibition

We previously showed using electron microscopy (EM) that constitutive mTORC1 activation due to loss of Tsc1 causes a significant enlargement of DA axon terminals, which is most pronounced in the SNc-innervated dorsolateral striatum (DLS) (Kosillo et al., 2019). In DA-Tsc1 KO mice, DA release was strongly reduced despite increased DA synthesis, suggesting that alterations in axon terminal structure may be an important determinant of DA releasability. To determine how suppression of mTORC1 or mTORC2 signaling affects the structural properties of DA axons, we developed a workflow for quantitatively assessing DA axon density and size. These parameters are challenging to measure accurately with conventional microscopy due to the high density of dopaminergic axons within the striatum. To overcome this, we employed protein-retention expansion microscopy (ProExM), whereby physical enlargement of the tissue effectively increases the resolution of light microscopy. We combined ProExM with light sheet fluorescence imaging and examined global DA axon architecture in DA-Raptor and DA-Rictor KO mice bred to the Ai9 tdTomato Cre reporter line (Madisen et al., 2010) (Figure 3a, Supplemental Figure 2a, Supplemental Video 1). To control for heterogeneous tdTomato expression along DA axon segments, across striatal sub-regions, and between genotypes, automated segmentation of DA axons from light sheet volumes was performed using a 3D convolutional neural network, TrailMap (Friedmann et al., 2020). We validated our DA axon segmentation pipeline in DA-Tsc1 KO mice, which have axon terminal hypertrophy as measured by EM (Kosillo et al., 2019). Consistent with the EM analysis, quantification of TrailMap-segmented images revealed an increase in the total axon volume of striatal DA-Tsc1 KO projections, which was due to an increase in both axonal density and the radius of individual axon segments (Supplemental Figure 2b-e, Supplemental Videos 2 and 3).

**Figure 3.**
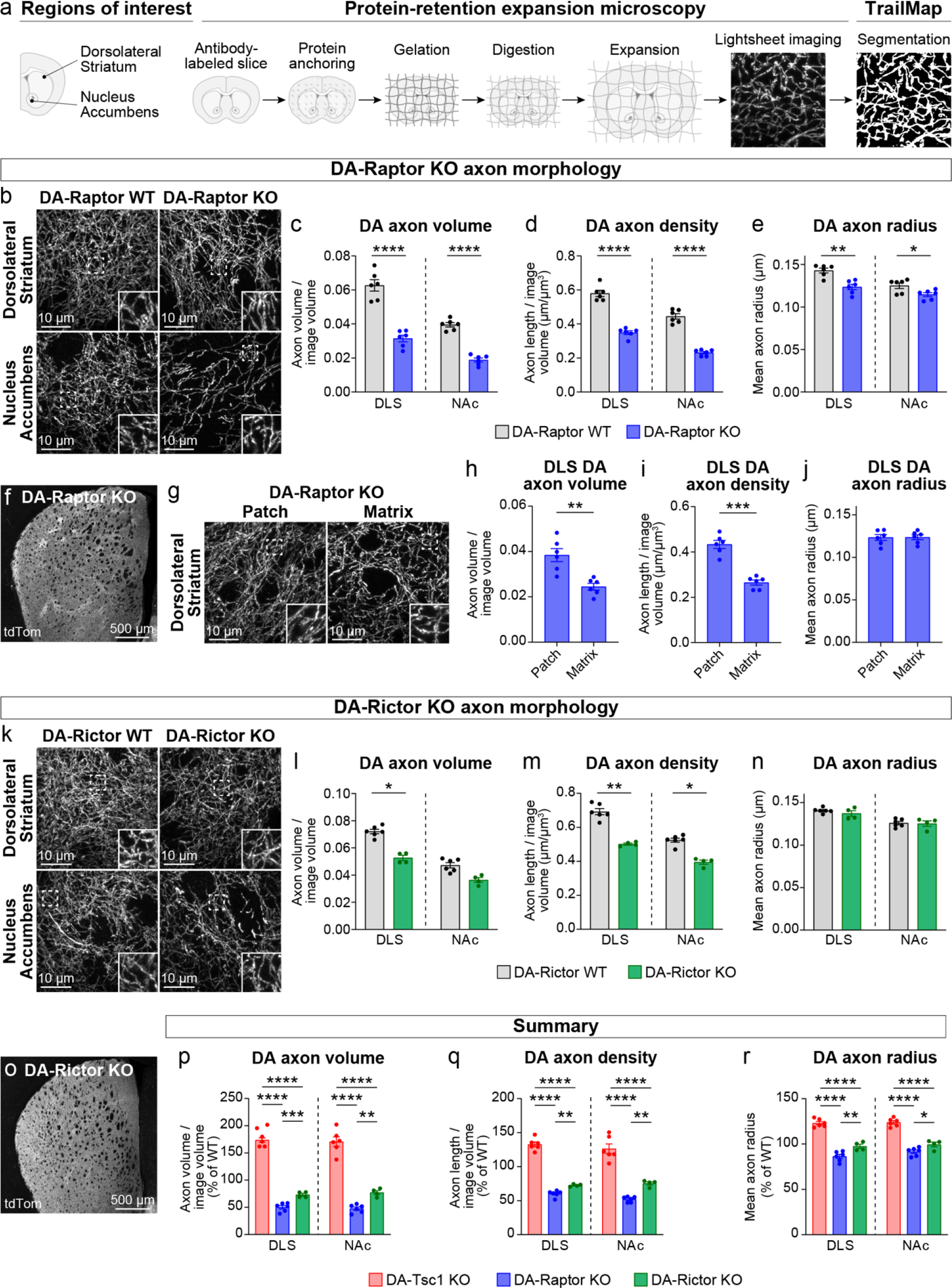
Dopamine axon density is reduced by inhibition of mTORC1 or mTORC2. **a)** Outline of the expansion microscopy and analysis workflow. Striatal slices containing tdTomato-labeled DA axons were labeled with an anti-RFP antibody and processed for protein-retention expansion microscopy (ProExM). Regions of interest containing dorsolateral striatum (DLS) or nucleus accumbens core (NAc) were imaged with light sheet fluorescence microscope and DA axons were segmented using TrailMap. **b)** Representative light sheet images of ProExM-processed striatal DA axons from DA-Raptor WT (left panels) and DA-Raptor KO (right panels) mice. Scale bars are normalized by expansion factor. Insets show high magnification images of boxed regions from the same panels. **c-e)** Mean ± SEM DA axon volume (**c**), density (**d**), and radius (**e**) in the DLS and NAc regions from DA-Raptor WT and DA-Raptor KO mice. n=6 slices from 3 mice per genotype (values are the average of 4 images per slice). Axon volume (**c**), ****p<0.0001 WT vs KO in DLS, ****p<0.0001 WT vs KO in NAc. Axon density (**d**), ****p<0.0001 WT vs KO in DLS, ****p<0.0001 WT vs KO in NAc. Axon radius (**e**), **p=0.0011 WT vs KO in DLS, *p=0.0279 WT vs KO in NAc. Welch’s unpaired t-tests. **f)** Representative confocal image of an unexpanded DA-Raptor KO striatal section, visualized by Cre-dependent tdTomato (tdTom). **g)** Representative light sheet images of ProExM-processed DLS patch and matrix regions from DA-Raptor KO mice showing tdTomato-labeled DA axons, scale bars=10 μm (normalized by expansion factor). Insets show higher magnification images of boxed regions from the same panels. **h-j)** Mean ± SEM DA axon volume (**h**), density (**i**), and radius (**j**) in the DLS patch and matrix compartments from DA-Raptor KO mice. Axon volume (**h**), **p=0.0019. Axon density (**i**), ***p=0.0003. Axon radius (**j**), p=0.7536. Paired t-tests. n=6 slices from 3 mice (values are the average of 4 images per slice). **k)** Representative light sheet images of ProExM-processed striatal DA axons from DA-Rictor WT (left panels) and DA-Rictor KO (right panels) mice. Scale bars are normalized by expansion factor. Insets show high magnification images of boxed regions from the same panels. **l-n)** Mean ± SEM DA axon volume (**l**), density (**m**), and radius (**n**) in the DLS and NAc regions from DA-Rictor WT and DA-Rictor KO mice. For DA-Rictor WT, n=6 slices from 3 mice. For DA-Rictor KO, n=4 slices from 2 mice (values are the average of 4 images per slice). Axon volume (**l**), *p=0.0351 WT vs KO in DLS, p=0.1164 WT vs KO in NAc. Axon density (**m**), **p=0.0068 WT vs KO in DLS, *p=0.0184 WT vs KO in NAc. Axon radius (**n**), p=0.8059 WT vs KO in DLS, p=0.6195 WT vs KO in NAc. Welch’s unpaired t-tests. **o)** Representative confocal image of an unexpanded DA-Rictor KO striatal section, visualized by Cre-dependent tdTomato. **p-r)** Summary data across all genotypes examined. Mean ± SEM DA axon volume (**p**), density (**q**), and radius (**r**) in the DLS and NAc regions from DA-Tsc1 KO, DA-Raptor KO, and DA-Rictor KO mice expressed as a percentage of WT values for each genotype. Axon volume (**p**), DLS: p<0.0001, one-way ANOVA; Holm-Sidak’s multiple comparisons, ****p<0.0001 DA-Tsc1 KO vs DA-Raptor KO, ****p<0.0001 DA-Tsc1 KO vs DA-Rictor KO, ***p=0.0010 DA-Raptor KO vs DA-Rictor KO. NAc: p<0.0001, one-way ANOVA; Holm-Sidak’s multiple comparisons, ****p<0.0001 DA-Tsc1 KO vs DA-Raptor KO, ****p<0.0001 DA-Tsc1 KO vs DA-Rictor KO, **p=0.0076 DA-Raptor KO vs DA-Rictor KO. Axon density (**q**), DLS: p<0.0001, one-way ANOVA; Holm-Sidak’s multiple comparisons, ****p<0.0001 DA-Tsc1 KO vs DA-Raptor KO, ****p<0.0001 DA-Tsc1 KO vs DA-Rictor KO, **p<0.0087 DA-Raptor KO vs DA-Rictor KO. NAc: p<0.0001, one-way ANOVA; Holm-Sidak’s multiple comparisons, ****p<0.0001 DA-Tsc1 KO vs DA-Raptor KO, ****p<0.0001 DA-Tsc1 KO vs DA-Rictor KO, **p=0.0070 DA-Raptor KO vs DA-Rictor KO. Axon radius (**r**), DLS: p<0.0001, one-way ANOVA; Holm-Sidak’s multiple comparisons, ****p<0.0001 DA-Tsc1 KO vs DA-Raptor KO, ****p<0.0001 DA-Tsc1 KO vs DA-Rictor KO, **p=0.0038 DA-Raptor KO vs DA-Rictor KO. NAc: p<0.0001, one-way ANOVA; Holm-Sidak’s multiple comparisons, ****p<0.0001 DA-Tsc1 KO vs DA-Raptor KO, ****p<0.0001 DA-Tsc1 KO vs DA-Rictor KO, *p=0.0210 DA-Raptor KO vs DA-Rictor KO. See also (Supplemental Figure 2) and Supplemental Videos 1-7.

We applied the validated ProExM-TrailMap pipeline to striatal slices from DA-Raptor-KO;Ai9 mice and found that, consistent with their somatodendritic hypotrophy, DA-Raptor KO axons in the DLS and NAc core exhibited a significant decrease in axon volume driven by a reduction in both axon density and radius (Figure 3b-e, Supplemental Videos 4 and 5). These phenotypes are opposite to what we observed in DA-Tsc1 KO slices (see Supplementary Figure 2b-e), demonstrating that mTORC1 signaling bi-directionally controls striatal DA axon density and size. We further noted that the striatal tdTomato fluorescence intensity in DA-Raptor KO mice showed regional heterogeneity, appearing as a patchwork pattern, which was particularly apparent within the DLS (Figure 3f). The striatal “patches” with brighter tdTomato signal in DA-Raptor KO mice aligned with striatal patches (also called striosomes) defined by high mu opioid receptor expression (Supplemental Figure S2f). The total area of patches was similar between genotypes, suggesting that the global patch-matrix striatal compartmentalization was intact (Supplemental Figure S2g). However, the ratio of tdTomato fluorescence in patch versus matrix regions was significantly higher in DA-Raptor KO slices compared to WT (Supplemental Figure S2h). We therefore separately imaged axons within the patches and the adjacent matrix in DLS slices from DA-Raptor KO mice (Figure 3g and Supplemental Video 4). The volume of DA axons within matrix regions was significantly reduced compared to patch regions, which was due to a selective reduction in axon density, as axon radius was similar across both compartments (Figure 3h-j). These results indicate that mTORC1 inhibition reduces the size and density of DA axons throughout the striatum, with a more pronounced deficit in axonal innervation of the matrix compartment.

We performed ProExM-TrailMap on striatal slices from DA-Rictor-KO;Ai9 mice and found that DA-Rictor KO projections in the DLS and NAc core exhibited a small decrease in axon volume that was driven by a change in axon density but not radius (Figure 3k-n, Supplemental Videos 6 and 7). Similar to WT mice, DA-Rictor KO axons showed homogenous innervation of the striatum, with no discernable regional differences in the tdTomato signal (Figure 3o). Taken together, the analysis of DA axons using ProExM-TrailMap demonstrates that constitutive mTORC1 suppression leads to DA axonal hypotrophy characterized by a concurrent reduction in axon density and radius. In contrast, mTORC2 suppression results in a more moderate decrease in axonal volume due to a selective reduction in axon density (Figure 3p-r).

### Electrophysiological properties of DA neurons are differentially altered by mTORC1 or mTORC2 suppression

Given the significant changes in DA neuron morphology observed in DA-Raptor KO mice, and to a lesser extent DA-Rictor KO mice, we performed whole-cell recordings to determine how these structural changes affected intrinsic membrane properties and excitability. Consistent with the somatodendritic and axonal hypotrophy of DA-Raptor KO neurons, we observed a large decrease in membrane capacitance and increase in membrane resistance in SNc DA-Raptor KO cells compared to controls (Figure 4a-b), with no changes to the resting membrane potential (Supplemental Table 1). In VTA neurons, deletion of *Rptor* decreased capacitance (Figure 4f) but did not significantly alter membrane resistance (Figure 4g) or membrane potential (Supplemental Table 1) compared to WT cells.

**Figure 4.**
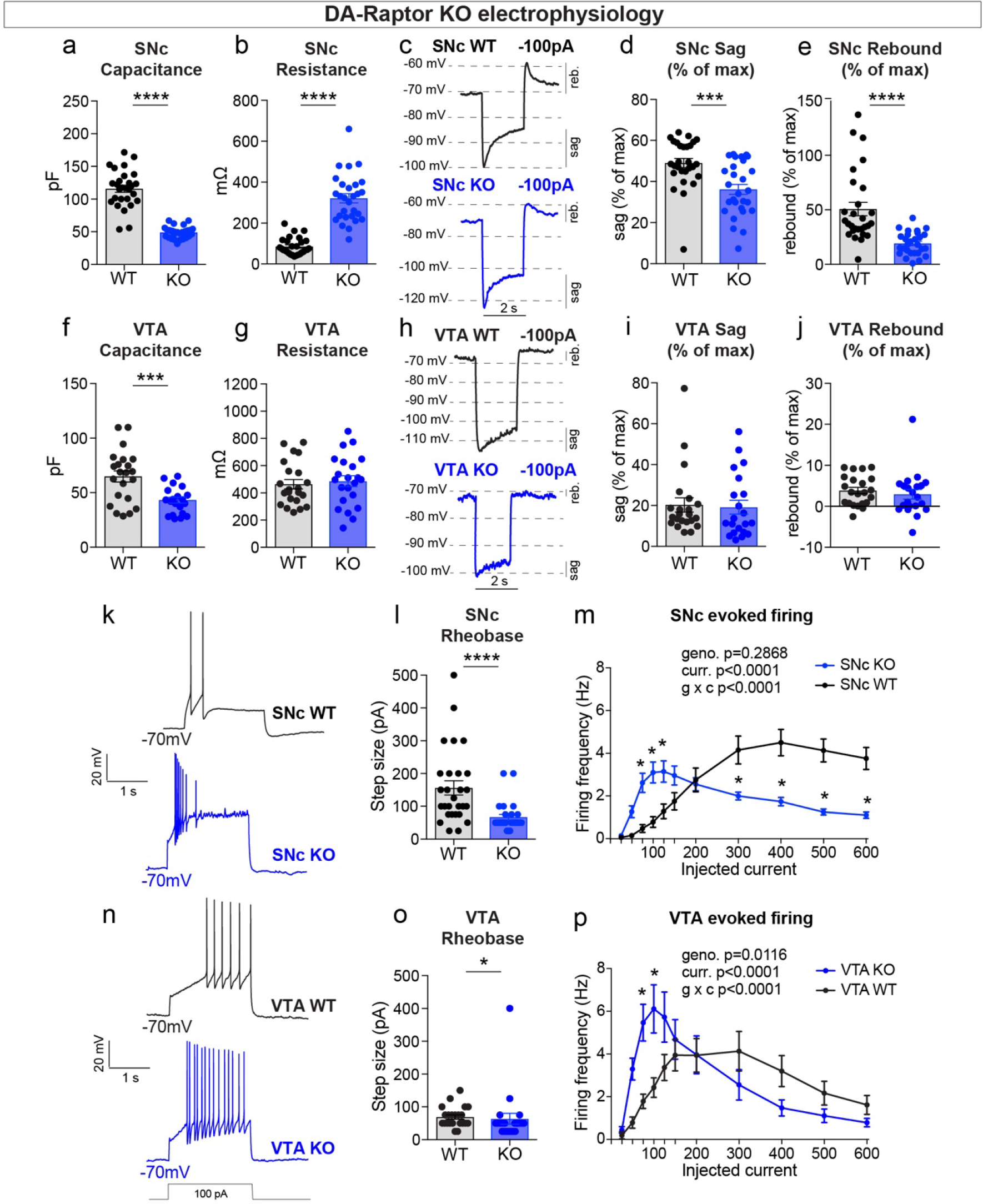
mTORC1 suppression increases the excitability of SNc and VTA DA neurons. **a,b)** Mean ± SEM membrane capacitance (**a**) and membrane resistance (**b**) of SNc DA neurons. DA-Raptor WT in black: n=28 neurons from eight mice, DA-Raptor KO in blue: n=28 neurons from six mice. Capacitance (**a**), ****p<0.0001, unpaired two-tailed t-test. Resistance (**b**), ****p<0.0001, Mann–Whitney test. **c)** Example current-clamp recordings from SNc DA neurons of the indicated genotypes in response to a −100 pA current step. reb.=rebound. **d,e)** Mean ± SEM sag (**d**) and rebound (**e**) amplitude expressed as a percentage of the maximum hyperpolarization from baseline in SNc DA neurons. DA-Raptor WT in black: n=28 neurons from six mice, DA-Raptor KO in blue: n=28 neurons from six mice. Sag (**d**), ***p=0.0003. Rebound (**e**), ****p<0.0001. Mann–Whitney tests. **f,g)** Mean ± SEM membrane capacitance (**f**) and membrane resistance (**g**) of VTA DA neurons. DA-Raptor WT in black: n=22 neurons from eight mice, DA-Raptor KO in blue: n=22 neurons from six mice. Capacitance (**f**), ***p=0.0005. Resistance (**g**), p=0.6769. Unpaired two-tailed t-tests. **h)** Example current-clamp recordings from VTA DA neurons of the indicated genotypes in response to a −100 pA current step. **i,j)** Mean ± SEM sag (**i**) and rebound (**j**) amplitude expressed as a percentage of the maximum hyperpolarization from baseline in VTA DA neurons. DA-Raptor WT in black: n=22 neurons from six mice, DA-Raptor KO in blue: n=21 neurons from six mice. Sag (**i**), p=0.3660. Rebound (**j**), p=0.2513. Mann–Whitney tests. **k)** Examples of action potential firing elicited with a +100 pA current step in SNc DA neurons of the indicated genotypes. **l)** Mean ± SEM rheobase of SNc DA neurons calculated as the current at which action potentials were first elicited. DA-Raptor WT in black: n=28 neurons from eight mice, DA-Raptor KO in blue: n=27 neurons from six mice, ****p < 0.0001, Mann–Whitney test. **m)** Input-output curves showing the firing frequency of SNc DA neurons in response to positive current steps of increasing amplitude. Data are displayed as mean ± SEM. DA-Raptor WT in black: n=28 neurons from eight mice, DA-Raptor KO in blue: n=28 neurons from six mice. Two-way ANOVA p values are shown. p_25pA_>0.9999, p_50pA_=0.3649, *p_75pA_=0.0009, *p_100pA_=0.0002 *p_125pA_=0.0069, p_150pA_=0.2497, p_200pA_>0.9999, *p_300pA_=0.0009, *p_400pA-600pA_<0.0001, Sidak’s multiple comparisons tests. **n)** Examples of action potential firing elicited with a +100 pA current step in VTA DA neurons of the indicated genotypes. **o)** Mean ± SEM rheobase of VTA DA neurons calculated as the current at which action potentials were first elicited. DA-Raptor WT in black: n=22 neurons from eight mice, DA-Raptor KO in blue: n=22 neurons from six mice, *p=0.0190, Mann–Whitney test. **p)** Input-output curves showing the firing frequency of VTA DA neurons in response to current steps of increasing amplitude. Data are displayed as mean ± SEM. DA-Raptor WT in black: n=22 neurons from eight mice, DA-Raptor KO in blue: n=22 neurons from six mice. Two-way ANOVA p values are shown. p_25pA_>0.9999, p_50pA_=0.0902, *p_75pA_=0.0014, *p_100pA_=0.0014, p_125pA_=0.1398, p_150pA_=0.9983, p_200pA_>0.9999, p_300pA_=0.6959, p_400pA_=0.5691, p_500pA_=0.9718, p_600pA_=0.9963, Sidak’s multiple comparisons tests. For all bar graphs, dots represent values for individual neurons. See also Supplemental Table 1.

Upon hyperpolarization, SNc DA neurons show a prominent sag response driven by hyperpolarization-activated (I_h_) current and a rebound response that depends on I_h_, type-A potassium current (I_A_) and T-type calcium channels (Amendola et al., 2012; Evans et al., 2017). This sag-rebound signature distinguishes SNc DA neurons from the neighboring VTA DA cells (Lammel et al., 2008). In response to a negative current step (−100 pA), we observed a significant reduction in both the sag and rebound responses of DA-Raptor KO SNc neurons compared to WT controls (Figure 4c-e). We plotted this as a percentage of the maximum hyperpolarization from baseline to account for the increased membrane resistance of DA-Raptor KO neurons, which influenced their degree of hyperpolarization (Figure 4d-e and Supplemental Table 1). In contrast to SNc neurons, mTORC1 suppression in VTA neurons had no impact on responses to hyperpolarizing current (Figure 4h-j).

We next examined responses to depolarizing current steps in DA-Raptor KO neurons to assess their intrinsic excitability (Figure 4k-p). We found a significant decrease in the minimum current required to evoke an action potential (rheobase) in DA-Raptor KO neurons compared to WT (Figure 4l,o). The input-output relationship of SNc and VTA DA-Rptor KO neurons to current steps of increasing amplitude was also altered such that they had increased firing frequency compared to WT cells at currents <150 pA, but exhibited substantial depolarization block and reduced firing rates at currents >300 pA (Figure 4m,p). Thus, constitutive mTORC1 inhibition increased the intrinsic excitability of both SNc and VTA neurons at lower current amplitudes, while firing was impaired in response to large depolarizing currents.

We measured the properties of DA-Rictor KO neurons and found that there was a small but significant decrease in capacitance for both SNc and VTA neurons (Figure 5a,f), consistent with their modest decrease in soma size (see Figure 2). VTA, but not SNc, DA-Rictor KO neurons also exhibited a small but significant increase in membrane resistance (Figure 5b,g), with no change in resting membrane potential (Supplemental Table 1). In contrast to loss of *Rptor*, deletion of *Rictor* did not affect sag-rebound responses in either SNc (Figure 5c-e) or VTA neurons (Figure 5h-j). We assessed intrinsic excitability and found that DA-Rictor KO SNc neurons had no change in rheobase and only a small shift in the input-output curve, which trended towards decreased excitability with higher current injections (Figure 5k-m). In contrast, DA-Rictor KO VTA neurons exhibited a significant decrease in rheobase and increased firing in response to current steps between 75 and 200 pA (Figure 5n-p). Together, these data show that mTORC1 suppression alters multiple aspects of DA neuron physiology in both SNc and VTA DA neurons, while mTORC2 inhibition primarily affects the excitability of VTA DA neurons.

**Figure 5.**
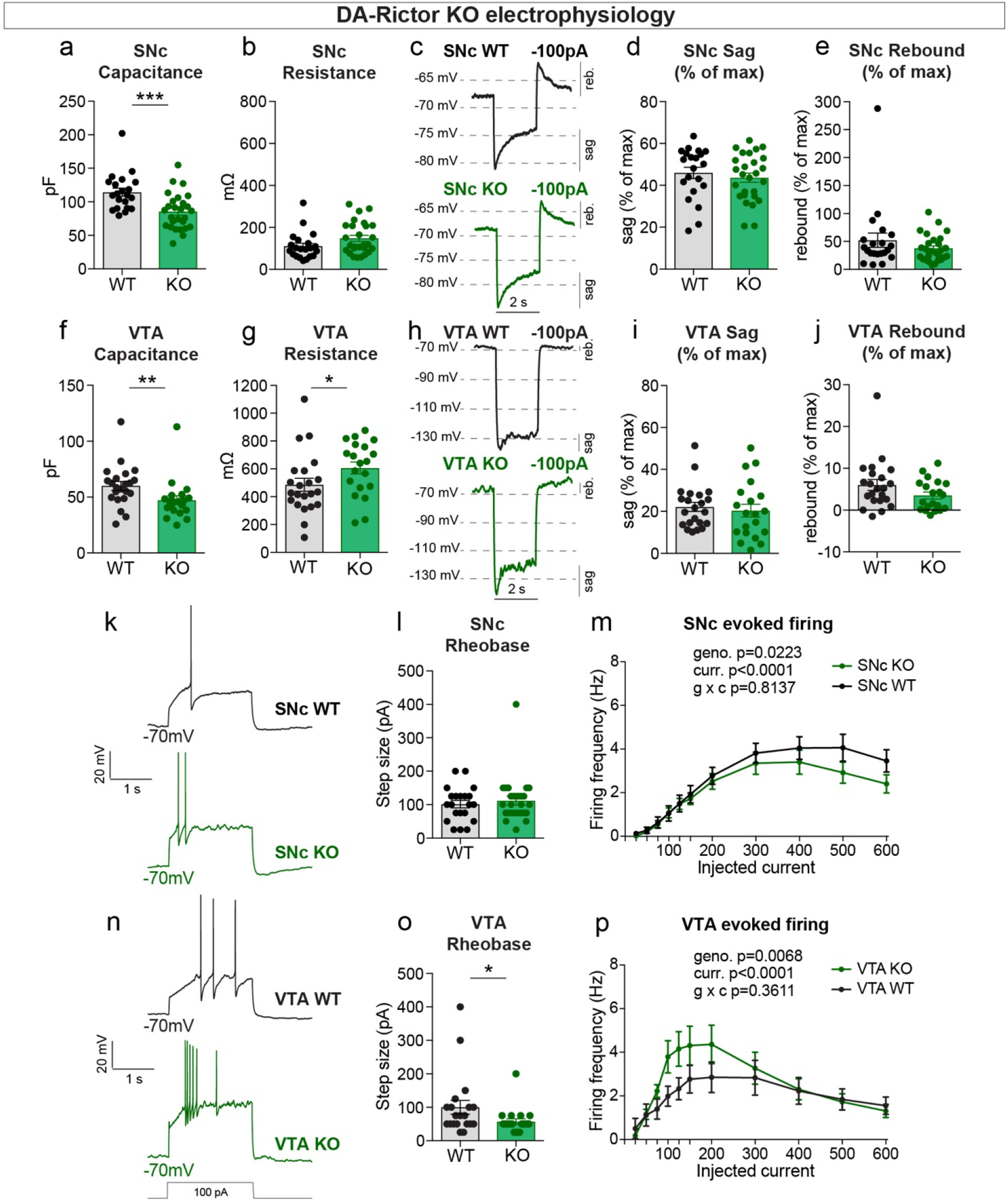
mTORC2 inhibition increases the excitability of VTA DA neurons. **a,b)** Mean ± SEM membrane capacitance (**a**) and membrane resistance (**b**) of SNc DA neurons. DA-Rictor WT in black: n=21 neurons from four mice, DA-Rictor KO in green: n=27 neurons from eight mice. Capacitance (**a**), ***p=0.0006. Resistance (**b**), p=0.0772 (**b**). Mann–Whitney tests. **c)** Example current-clamp recordings from SNc DA neurons of the indicated genotypes in response to a −100 pA current step. **d,e)** Mean ± SEM sag (**d**) and rebound (**e**) amplitude expressed as a percentage of the maximum hyperpolarization from baseline in SNc DA neurons. DA-Rictor WT in black: n = 21 neurons from four mice, DA-Rictor KO in green: n = 27 neurons from eight mice. Sag (**d**), p=0.5006, two-tailed t-test. Rebound (**e**), p=0.4212, Mann–Whitney test. **f,g)** Mean ± SEM membrane capacitance (**f**) and membrane resistance (**g**) of VTA DA neurons. DA-Rictor WT in black: n=22 neurons from four mice, DA-Rictor KO in green: n=20 neurons from six mice. Capacitance (**f**), **p=0.0014. Resistance (**g**), *p=0.0418. Mann-Whitney tests. **h)** Example current-clamp recordings from VTA DA neurons of the indicated genotypes in response to a −100 pA current step. **i,j)** Mean ± SEM sag (**i**) and rebound (**j**) amplitude expressed as a percentage of the maximum hyperpolarization from baseline in VTA DA neurons. DA-Rictor WT in black: n=22 neurons from four mice, DA-Rictor KO in green: n=20 neurons from six mice. Sag (**i**), p=0.3880. Rebound (**j**), p=0.1638. Mann–Whitney tests. **k)** Examples of action potential firing elicited with a +100 pA current step in SNc DA neurons of the indicated genotypes. **l)** Mean ± SEM rheobase of SNc DA neurons calculated as the current at which action potentials were first elicited. DA-Rictor WT in black: n=21 neurons from four mice, DA-Rictor KO in green: n=27 neurons from eight mice, p=0.7414, Mann–Whitney test. **m)** Input-output curves showing the firing frequency of SNc DA neurons in response to current steps of increasing amplitude. Data are displayed as mean ± SEM. DA-Rictor WT in black: n = 21 neurons from four mice, DA-Rictor KO in green: n = 27 neurons from eight mice. Two-way ANOVA p values are shown. p_25pA_>0.9999, p_50pA_>0.9999, p_75pA_>0.9999, p_100pA_>0.9999, p_125pA_>0.9999, p_150pA_>0.9999, p_200pA_>0.9999, p_300pA_=0.9948, p_400pA_=0.9341, p_500pA_=0.2913, p_600pA_=0.4672, Sidak’s multiple comparisons tests. **n)** Examples of action potential firing elicited with a +100 pA current step in VTA DA neurons of the indicated genotypes. **o)** Mean ± SEM rheobase of VTA DA neurons calculated as the current at which action potentials were first elicited. DA-Rictor WT in black: n=20 neurons from four mice, DA-Rictor KO in green: n=20 neurons from six mice, *p=0.0299, Mann–Whitney test. **p)** Input-output curves showing the firing frequency of VTA DA neurons in response to current steps of increasing amplitude. Data are displayed as mean ± SEM. DA-Rictor WT in black: n=22 neurons from four mice, DA-Rictor KO in green: n=20 neurons from six mice. Two-way ANOVA p values are shown. p_25pA_>0.9999, p_50pA_>0.9999, p_75pA_=0.9842, p_100pA_=0.2640, p_125pA_=0.2515, p_150pA_=0.4871, p_200pA_=0.5233, p_300pA_>0.9999, p_400pA_>0.9999, p_500pA_>0.9999, p_600pA_>0.9999, Sidak’s multiple comparisons tests. For all bar graphs, dots represent values for individual neurons. See also Supplemental Table 1.

### Evoked striatal DA release is differentially impacted by mTORC1 or mTORC2 inhibition

To determine how DA release is impacted by inhibition of mTORC1 or mTORC2, we used fast scan cyclic voltammetry (FCV) to measure evoked DA release within seven striatal subregions in DA-Raptor or DA-Rictor KO mice. We examined multiple striatal subregions since these areas are differentially innervated by DA neuron subpopulations and participate in distinct circuits and behavioral functions (Haber, 2014; Kramer et al., 2018; Lerner et al., 2015; Ogawa and Watabe-Uchida, 2018). The dorsal striatum (sites 1-6, Figure 6a) is preferentially innervated by SNc DA neurons, while the ventral striatum, or NAc (site 7, Figure 6a), is primarily targeted by VTA DA cells. Constitutive mTORC1 suppression in DA-Raptor KO mice resulted in a profound impairment in electrically-evoked DA release across all regions sampled (Figure 6a-c, Supplemental Figure 3a-c). We observed a more than 60% reduction in peak evoked DA release with both single pulse (Figure 6b,c) and high frequency burst (Supplemental Figure S3b,c) stimulation across dorsal and ventral striatal sampling sites. This level of impairment in evoked DA release is generally consistent with that observed in the dorsal striatum of DA-Tsc1 KO mice, in which mTORC1 is hyperactive (Kosillo et al., 2019). Thus, impaired evoked DA release can occur either with constitutive mTORC1 activation or suppression. We compared the kinetics of DA re-uptake via DAT and found a significant reduction in re-uptake rate in the dorsal striatum (sites 1-6, Figure 6d,e) but not the NAc (site 7, Figure 6f,g) of DA-Raptor KO mice, suggesting a region-specific decrease in DAT expression and/or activity. The decreased DA re-uptake rate may serve a compensatory function to boost dopaminergic signaling in response to compromised DA release. Correspondingly, baseline DAT expression is higher in dorsal striatum compared to ventral striatum (Condon et al., 2019; Rice and Cragg, 2008; Sulzer et al., 2016), which presents an additional measure of control for adjusting DA levels via changes in DAT.

**Figure 6.**
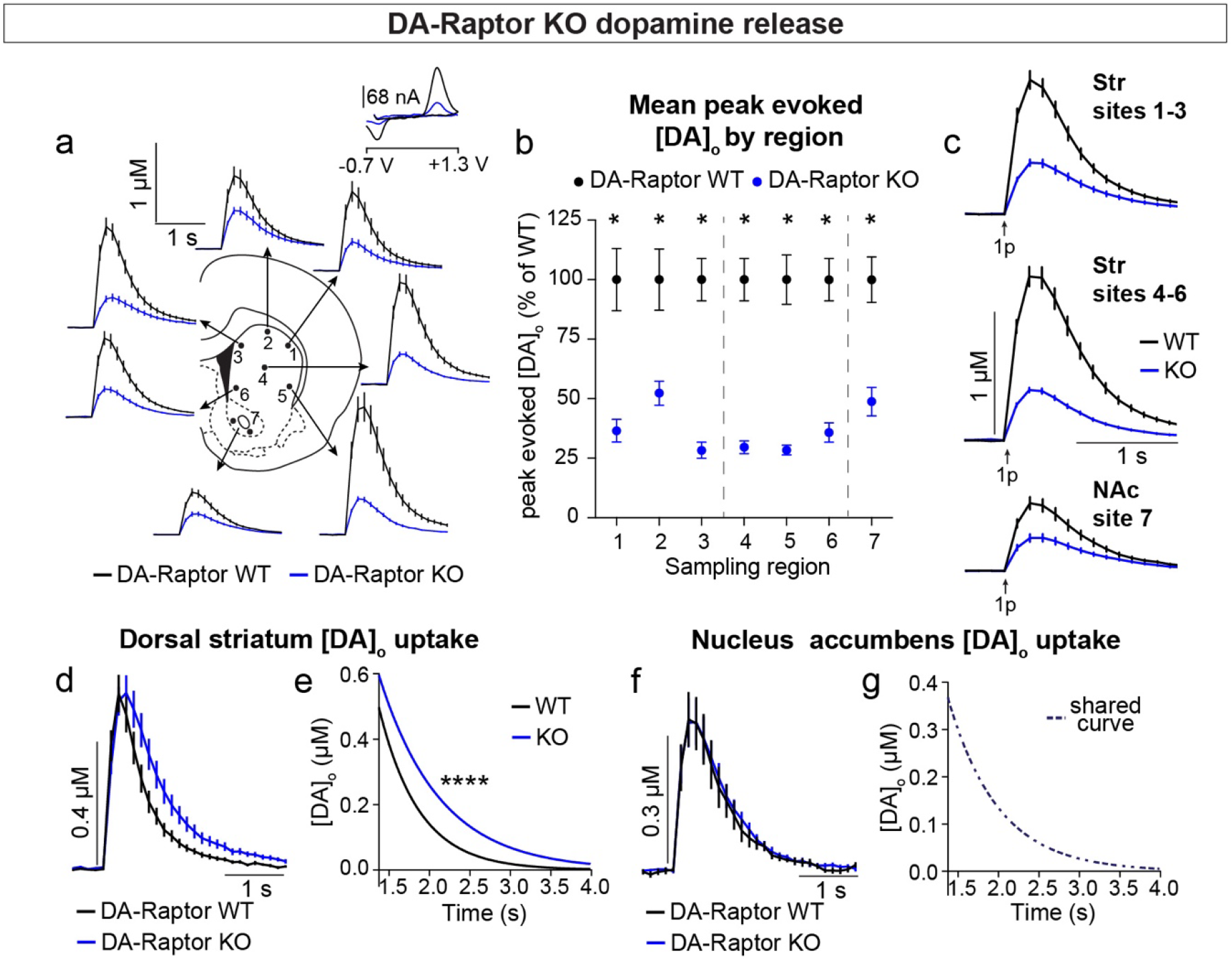
Deletion of *Rptor* reduces evoked DA release across all striatal regions. **a)** Mean ± SEM [DA]_o_ versus time evoked from different striatal subregions by a single electrical pulse. Traces are an average of n=20 (sites #1,2,4), n=18 (sites #3,6), n=19 (site #5) and n=34 (site #7) transients per sampling region from five mice per genotype. DA-Raptor WT in black, DA-Raptor KO in blue. Inset, typical cyclic voltammograms show characteristic DA waveform. **b)** Mean ± SEM peak [DA]_o_ by striatal subregion expressed as a percentage of WT (sampling region numbers correspond to the sites in panel **a**). n=20 transients (sites #1,2,4), n=18 transients (sites #3,6), n=19 transients (site #5) and n=34 transients (site #7) per sampling region from five mice per genotype. *p_1_=0.0002, *p_3_<0.0001, *p_7_<0.0001, Wilcoxon’s two-tailed t-tests; *p_2_=0.0047, *p_4_<0.0001, *p_5_<0.0001, *p_6_<0.0001, paired two-tailed t-tests. **c)** Mean ± SEM [DA]_o_ versus time averaged across all transients from three striatal territories, dorsal striatum (Str) (sites #1–3), central-ventral striatum (sites #4-6) and NAc core (site #7). Traces are an average of n=58 transients (sites #1-3), n=57 transients (sites #4-6) and n=34 transients (site #7) per sampling territory from five mice per genotype. Statistical comparisons for the peak evoked [DA]_o_ between genotypes by sub-region: ****p_Str 1-3_<0.0001, ****p_NAc 7_<0.0001, Wilcoxon’s two-tailed t-tests; ****p_Str 4-6_<0.0001, paired two-tailed t-tests. **d)** Mean ± SEM [DA]_o_ versus time from concentration- and site-matched FCV transients recorded in dorsal and central-ventral striatum (sites #1-6). DA-Raptor WT average of n=20 transients from five mice per genotype, DA-Raptor KO average of n=22 transients from five mice per genotype. **e)** Single-phase exponential decay curve-fit of the falling phase of concentration- and site- matched striatal DA transients (from panel **d**). X-axis starts 375 ms after stimulation. DA-Raptor WT average of n=20 transients from five mice per genotype, DA-Raptor KO average of n=22 transients from five mice per genotype. ****p<0.0001, least-squares curve-fit comparison. **f)** Mean ± SEM [DA]_o_ versus time from concentration- and site-matched FCV transients recorded in NAc core (site #7). DA-Raptor WT average of n=9 transients from five mice per genotype, DA-Raptor KO average of n=10 transients from five mice per genotype. **g)** Single-phase exponential decay curve-fit of the falling phase of concentration-matched NAc DA transients (from panel **f**). X-axis starts 375 ms after stimulation onset. DA-Raptor WT average of n=9 transients from five mice per genotype, DA-Raptor KO average of n=10 transients from five mice per genotype. p=0.4377, least-squares curve-fit comparison. See also (Supplemental Figure 3).

Constitutive mTORC2 suppression in DA-Rictor KO mice resulted in a different profile of striatal DA release. In DA-Rictor KO mice, we found a moderate reduction in peak evoked DA release specifically in the ventral half of the striatum and NAc (sites 4-7), while DA release in the dorsal-most regions (sites 1-3) was unchanged (Figure 7a-c and Supplemental Figure 3d-f). On average, we observed an approximately 25% reduction in peak evoked DA release both with single pulse (Figure 7b,c) and high frequency burst (Supplemental Figure S3e,f) stimulation at sampling sites 4-7, which receive proportionately denser innervation from VTA compared to SNc DA neurons (Chen et al., 2021). The observation that release impairments in DA-Rictor KO mice were selectively present in ventral regions is consistent with the more pronounced electrophysiological changes we found in VTA neurons compared to SNc neurons with *Rictor* deletion (see Figure 5). We observed no change in DA re-uptake kinetics in either the dorsal striatum (Figure 7d,e) or NAc (Figure 7f,g) of DA-Rictor KO slices.

**Figure 7.**
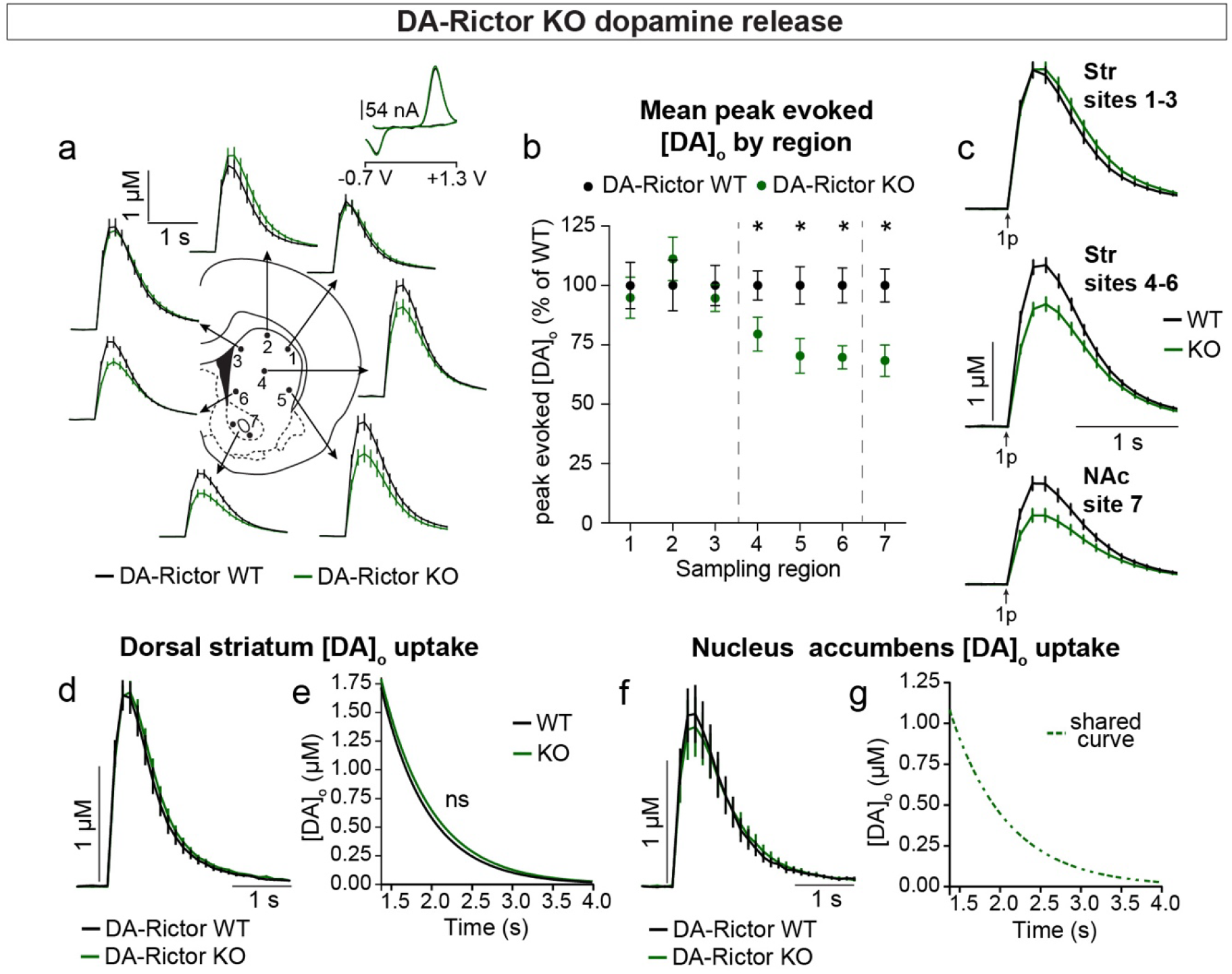
Deletion of *Rictor* reduces evoked DA release in central-ventral striatum and NAc. **a)** Mean ± SEM [DA]_o_ versus time evoked from different striatal subregions by a single electrical pulse. Traces are an average of n=20 (sites #1-6) or n=40 (site #7) transients per sampling region from five mice per genotype. DA-Rictor WT in black, DA-Rictor KO in green. Inset, typical cyclic voltammograms show characteristic DA waveform. **b)** Mean ± SEM peak [DA]_o_ by striatal subregion expressed as a percentage of WT (sampling region numbers correspond to the sites in panel **a**). n=20 (sites #1-6) or n=40 (site #7) transients per sampling region from five mice per genotype. p_1_=0.5185, p_2_=0.2858, p_3_=0.4538, *p_4_=0.0111, *p_5_=0.0161, *p_6_=0.0003, paired two-tailed t-tests; *p_7_<0.0001, Wilcoxon’s two-tailed t-test **c)** Mean ± SEM [DA]_o_ versus time averaged across all transients from three striatal territories, dorsal striatum (Str) (sites #1-3), central-ventral striatum (sites #4-6) and NAc core (site #7). Traces are an average of n=60 transients (sites #1-3, 4-6) or n=40 transients (site #7) per sampling territory from five mice per genotype. Statistical comparisons for the peak evoked [DA]_o_ between genotypes by sub-region: p_Str 1-3_=0.9672, paired two-tailed t-test; ****p_Str 4-6_<0.0001, ****p_NAc 7_<0.0001, Wilcoxon’s two-tailed t-tests. **d)** Mean ± SEM [DA]_o_ versus time from concentration- and site-matched FCV transients recorded in dorsal and central-ventral striatum (sites #1-6). DA-Rictor WT average of n=16 transients from five mice per genotype, DA-Rictor KO average of n=17 transients from five mice per genotype. **e)** Single-phase exponential decay curve-fit of the falling phase of concentration- and site- matched striatal DA transients (from panel **d**). X-axis starts 375 ms after stimulation. DA-Rictor WT average of n=16 transients from five mice per genotype, DA-Rictor KO average of n=17 transients from five mice per genotype. p=0.0594, least-squares curve-fit comparison. **f)** Mean ± SEM [DA]_o_ versus time from concentration- and site-matched FCV transients recorded in NAc core (site #7). DA-Rictor WT average of n=10 transients from five mice per genotype, DA- Rictor KO average of n=10 transients from five mice per genotype. **g)** Single-phase exponential decay curve-fit of the falling phase of concentration-matched NAc DA transients (from panel **f**). X-axis starts 375 ms after stimulation onset. DA-Rictor WT average of n=10 transients from five mice per genotype, DA-Rictor KO average of n=10 transients from five mice per genotype. p=0.8759, least-squares curve-fit comparison. See also Supplemental Figure 3.

Together these data demonstrate that constitutive mTORC1 suppression results in a ubiquitous impairment in evoked striatal DA release and a likely compensatory decrease in DA clearance rate in the dorsal striatum. mTORC2 inhibition causes a milder impairment in DA release, which specifically affects ventral regions preferentially targeted by VTA projections.

### Inhibition of mTORC1 but not mTORC2 impairs DA synthesis

To investigate whether alterations in DA synthesis or vesicle packaging could account for the release impairments in DA-Raptor or DA-Rictor KO mice, we harvested striatal tissue and measured protein levels of the vesicular monoamine transporter 2 (VMAT-2), which is responsible for vesicular loading of DA, and TH, which is the rate-limiting enzyme in DA synthesis. In striatal lysates from DA-Raptor KO mice, we found no change in VMAT-2 expression compared to WT controls, suggesting that vesicular DA loading is likely intact (Figure 8a,b). However, striatal lysates from DA-Raptor KO mice showed a significant downregulation of TH protein expression (Figure 8a,b), consistent with their impaired DA release. TH levels were also reduced in DA cell bodies in both the SNc (Figure 8d-f) and VTA (Figure 8g-i). Together, these data suggest reduced DA synthesis capacity resulting from constitutive mTORC1 suppression. To examine DA synthesis and content directly, we harvested dorsal and ventral striatal tissue and measured DA levels using high performance liquid chromatography (HPLC). We found a large reduction in the total tissue DA content across both regions in DA-Raptor KO mice compared to controls, while the ratio of DA to its primary metabolite DOPAC was unchanged (Figure 8j,k, see Supplemental Table 2 for raw values). This profile is indicative of reduced DA production in DA-Raptor KO mice, rather than increased DA turnover.

**Figure 8.**
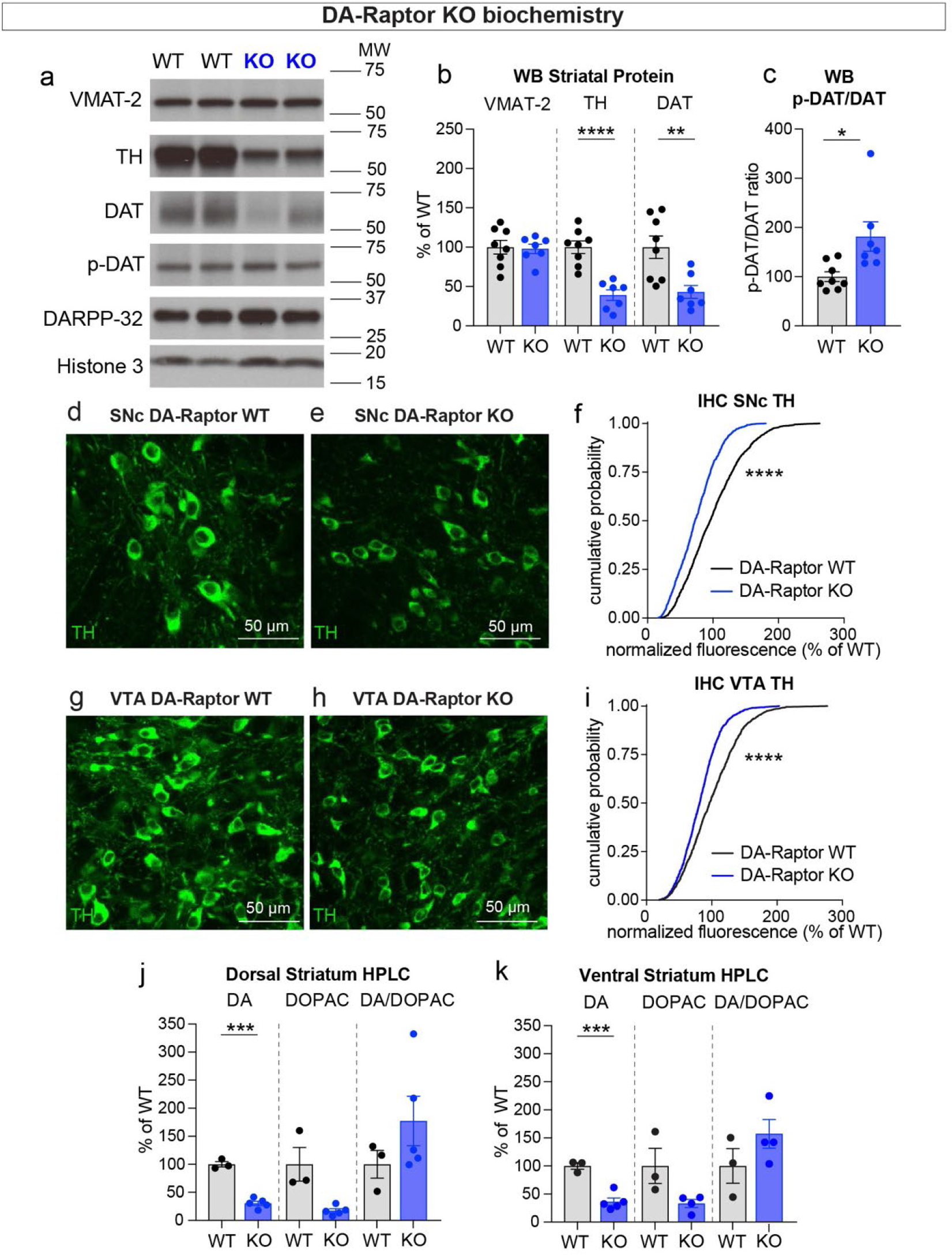
Deletion of *Rptor* reduces TH expression and DA synthesis. **a)** Striatal lysates were harvested from DA-Raptor WT or KO mice. Representative western blots for vesicular monoamine transporter-2 (VMAT-2), tyrosine hydroxylase (TH), dopamine active transporter (DAT), DAT phosphorylated at Thr53 (p-DAT), DARPP-32 and Histone 3. Two independent samples per genotype are shown. Observed molecular weight (MW) noted on the right. **b)** Mean ± SEM striatal protein content of VMAT-2, TH and DAT. Dots represent values for individual mice (averaged from two samples per mouse). n=8 DA-Raptor WT mice in black and n=7 DA-Raptor KO mice in blue. VMAT-2, p=0.8476; TH, ****p<0.0001; DAT, **p=0.0053, Welch’s two-tailed t-tests. **c)** Mean ± SEM ratio of DAT phosphorylated at Thr53 (p-DAT) to total DAT protein, *p=0.0340, Welch’s two-tailed t-test. **d,e)** Representative confocal images of SNc neurons from DA-Raptor WT (**d**) or DA-Raptor KO (**e**) mice immunostained for TH, scale bars=50 μm. **f)** Cumulative distribution of TH levels in SNc DA neurons. DA-Raptor WT in black: n=1024 neurons from three mice, DA-Raptor KO in blue: n=1045 neurons from three mice, ****p<0.0001, Kolmogorov–Smirnov test. **g,h)** Representative confocal images of VTA neurons from DA-Raptor WT (**g**) or DA-Raptor KO (**h**) mice immunostained for TH, scale bars=50 μm. **i)** Cumulative distribution of TH levels in VTA DA neurons. DA-Raptor WT in black: n=1389 neurons from three mice, DA-Raptor KO in blue: n=1526 neurons from three mice, ****p<0.0001, Kolmogorov–Smirnov test. **j,k)** Mean ± SEM total tissue content of DA, 3,4-dihydroxyphenylacetic acid (DOPAC) and the DOPAC/DA ratio per mouse assessed by HPLC from tissue punches from dorsal striatum (**j**) or ventral striatum (**k**). Dots represent values for individual mice (averaged from two samples per mouse). n=3 DA-Raptor WT mice in black and n=5 DA-Raptor KO mice in blue (n=4 DA-Raptor KO mice in blue for ventral striatum DOPAC). Dorsal striatum DA, ***p=0.0002, DOPAC, p=0.1084, DA/DOPAC ratio, p=0.1790, Welch’s two-tailed t-tests. Ventral striatum DA, ***p=0.0005, DOPAC, p=0.1599, DA/DOPAC ratio, p=0.2190, Welch’s two-tailed t-tests. See Supplemental Table 2 for the raw HPLC measurement values.

Given the altered re-uptake kinetics observed in DA-Raptor KO mice in the FCV experiments, we examined striatal protein levels of DAT and phosphorylated DAT at Thr53 (p-DAT), which promotes transporter function (Foster et al., 2012). Consistent with the decreased re-uptake rate observed by FCV (see Figure 6d,e), we found reduced total DAT levels in the striatum of DA-Raptor KO mice (Figure 8a,b). The ratio of p-DAT to total DAT was increased (Figure 8c), suggesting that the remaining DAT is functional.

Together these data show that constitutive mTORC1 suppression leads to downregulation of TH levels in both the somatic and axonal compartments, resulting in reduced DA synthesis capacity and tissue content. Furthermore, DA-Raptor KO mice have reduced axonal DAT expression, which may be a compensatory change to prolong DA actions in response to limited neurotransmitter availability and release.

We next investigated whether changes in DA synthesis or packaging might be responsible for the altered DA release in DA-Rictor KO mice. Similar to *Rptor* deletion, loss of *Rictor* did not affect expression of VMAT-2 within the striatum (Figure 9a,b). However, in contrast to mTORC1 inhibition, mTORC2 suppression did not affect striatal TH, DAT, or p-DAT levels (Figure 9a-c). In the midbrain, we found a small shift in the cumulative distribution of TH levels in individual SNc neurons in DA-Rictor KO mice compared to controls (Figure 9d-f). However, the average somatic TH level per mouse was not different between genotypes (WT, 100.00% +/- 2.50; KO, 101.10% +/- 6.58, p=0.9905, Welch’s t-test, normalized values). In VTA neurons, deletion of *Rictor* led to a small but significant decrease in somatic TH levels (Figure 9g-i). This small reduction in TH did not translate into significant changes in the tissue content of DA or DOPAC in either the dorsal or ventral striatum of DA-Rictor KO mice (Figure 9j,k, see Supplemental Table 2 for raw values). Together these data show that reduced DA availability alone is unlikely to account for the DA release deficits in DA-Rictor KO mice.

**Figure 9.**
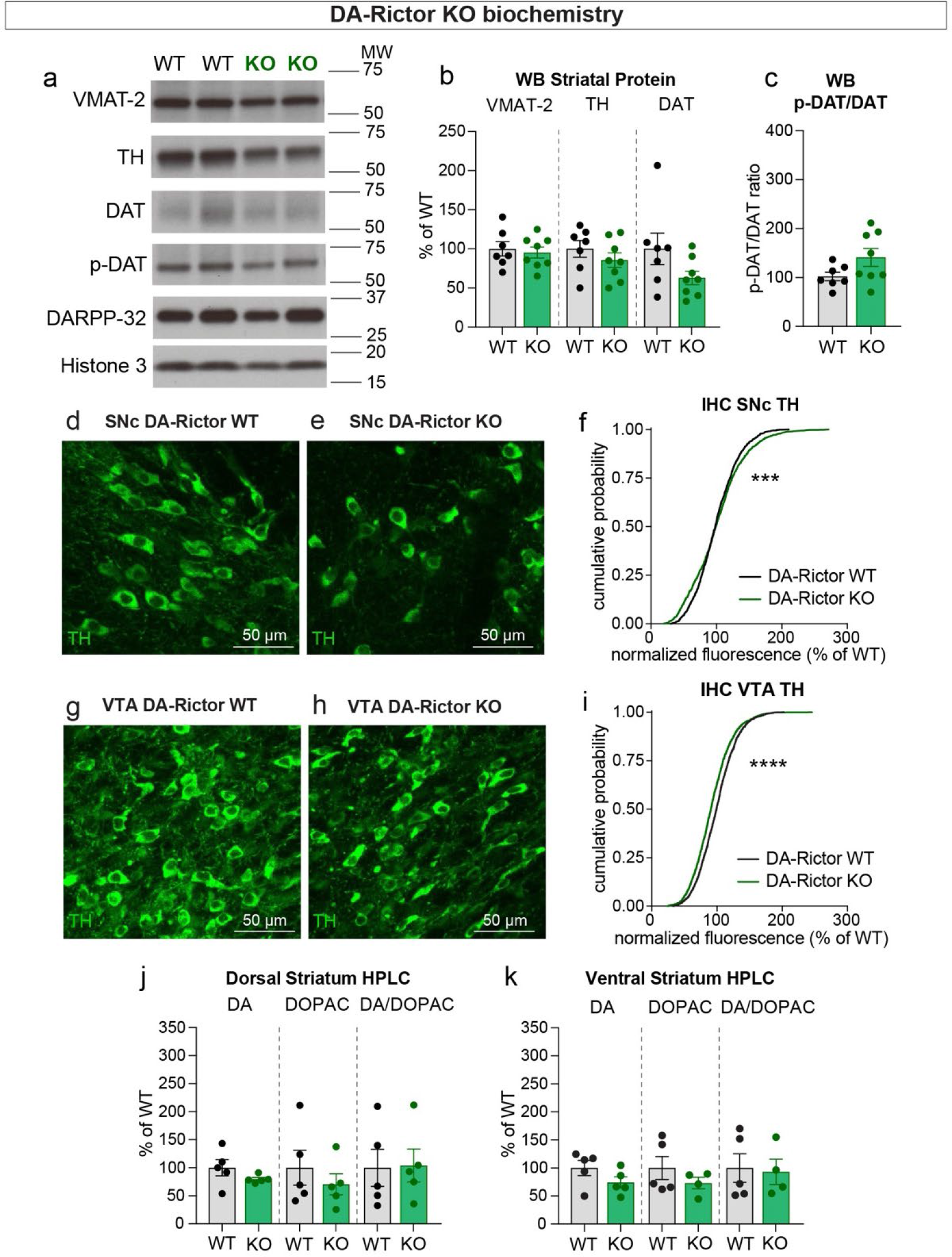
Deletion of *Rictor* does not alter striatal DA synthesis. **a)** Striatal lysates were harvested from DA-Rictor WT or KO mice. Representative western blots for vesicular monoamine transporter-2 (VMAT-2), tyrosine hydroxylase (TH), dopamine active transporter (DAT), DAT phosphorylated at Thr53 (p-DAT), DARPP-32 and Histone 3. Observed molecular weight (MW) noted on the right. **b)** Mean ± SEM striatal protein content of VMAT-2, TH and DAT. Dots represent values for individual mice (averaged from two samples per mouse). n=7 DA-Rictor WT mice in black and n=8 DA-Rictor KO mice in green. VMAT-2, p=0.6882; TH, p=0.3281; DAT, p=0.1282 Welch’s two-tailed t-tests. **c)** Mean ± SEM ratio of DAT phosphorylated at Thr53 (p-DAT) to total DAT protein, p=0.0838, Welch’s two-tailed t-test. **d,e)** Representative confocal images of SNc neurons from DA-Rictor WT (**d**) or DA-Rictor KO (**e**) mice immunostained for TH, scale bars=50 μm. **f)** Cumulative distribution of TH levels in SNc DA neurons. DA-Rictor WT in black: n=1280 neurons from three mice, DA-Rictor KO in green: n=1550 neurons from four mice, ***p=0.0009, Kolmogorov–Smirnov test. **g,h)** Representative confocal images of VTA neurons from DA-Rictor WT (**g**) or DA-Rictor KO (**h**) mice immunostained for TH, scale bars=50 μm. **i)** Cumulative distribution of TH levels in VTA DA neurons. DA-Rictor WT in black: n=1968 neurons from three mice, DA-Rictor KO in green: n=2370 neurons from four mice, ****p<0.0001, Kolmogorov–Smirnov test. **j,k)** Mean ± SEM total tissue content of DA, 3,4-dihydroxyphenylacetic acid (DOPAC) and the DOPAC/DA ratio per mouse assessed by HPLC from tissue punches from dorsal striatum (**j**) or ventral striatum (**k**). Dots represent values for individual mice (averaged from two samples per mouse). n=5 DA-Rictor WT mice in black and n=5 DA-Rictor KO mice in green (n=4 DA-Rictor KO mice in green for ventral striatum DOPAC). Dorsal striatum DA, p=0.2356, DOPAC, p=0.4438, DA/DOPAC ratio, p=0.9289, Welch’s two-tailed t-tests. Ventral striatum DA, p=0.1648, DOPAC, p=0.2896, DA/DOPAC ratio, p=0.8455, Welch’s two-tailed t-tests. See Supplemental Table 2 for the raw HPLC measurement values.

### Concurrent deletion of Rptor and Rictor exacerbates deficits in DA production, release and re-uptake

Given our observation that deletion of *Rictor* led to decreased p-S6 levels in DA neurons and that DA-Rictor KO mice shared several phenotypes with DA-Raptor KO mice, we asked whether the effects of *Rictor* deletion were mediated by mTORC1 suppression or whether mTORC2 might act via independent mechanisms. To test this, we generated double KO mice in which both *Rptor* and *Rictor* were selectively deleted from DA neurons (*Rptor^fl/fl^;Rictor^f/fl^;DAT- Cre^wt/+^;Ai9^+/-^*, Figure 10a-c). We compared mice with deletion of both *Rptor* and *Rictor* (DA-Raptor KO;Rictor KO, “Double KO”) to mice with deletion of *Rptor* alone (DA-Raptor KO;Rictor WT). With this strategy we could investigate whether loss of Rictor induced further changes in DA neuron properties in addition to those caused by Raptor loss or whether the changes observed in DA-Rictor KO neurons were due to reduced mTORC1 signaling. We found that loss of *Rptor* caused a maximal suppression of p-S6 levels in both SNc and VTA DA neurons, as concurrent deletion of *Rictor* did not further decrease p-S6 at our level of detection (Figure 10b-f, i-k). This suggests that the reduction in p-S6 in DA-Rictor KO mice (see Figure 2i,k) was likely due to suppression of mTORC1 activity. However, compared to *Rptor* deletion alone, concurrent deletion of *Rictor* and *Rptor* further reduced soma size and TH expression in both SNc (Figure 10g,h) and VTA (Figure 10l,m) neurons.

**Figure 10.**
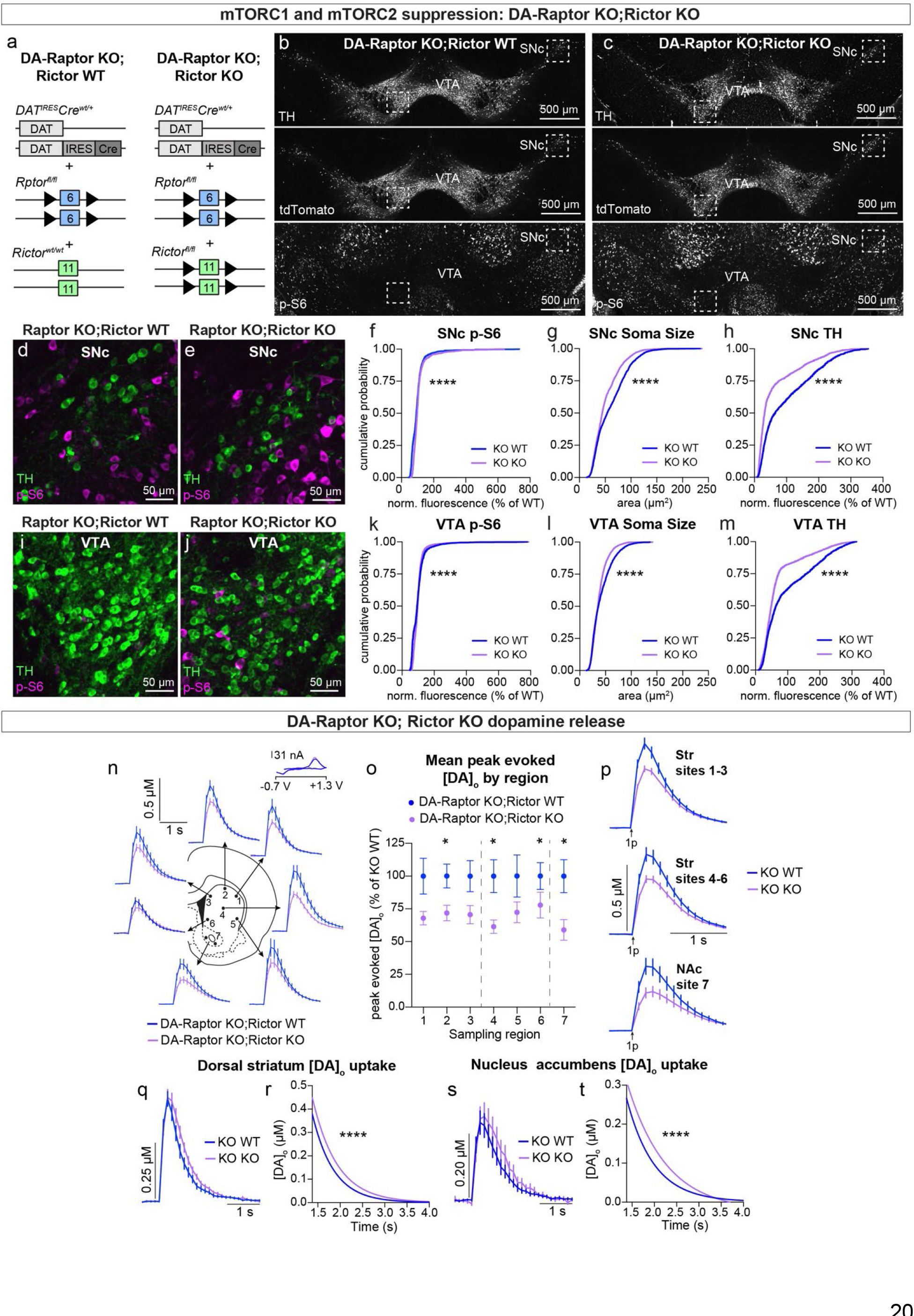
Double knock-out of *Rptor* and *Rictor* strongly impairs DA synthesis, release, and re-uptake. **a)** Schematic of the genetic strategy to delete either *Rptor* alone or *Rptor* and *Rictor* genes selectively from DA neurons. Numbered boxes represent exons and triangles represent loxP sites. **b,c)** Representative confocal images of midbrain sections from DA-Raptor KO;Rictor WT (**b**) and DA-Raptor KO;Rictor KO (**c**) mice with DA neurons visualized by Cre-dependent tdTomato expression and tyrosine hydroxylase (TH) immunostaining. Bottom panels show p-S6 (Ser240/244) immunostaining. Scale bars=500 μm. **d,e)** Higher magnification merged images of the boxed SNc regions in panels b and c, showing TH (green) and p-S6 (magenta), scale bars=50 μm. **f-h)** Cumulative distributions of SNc DA neuron p-S6 levels (**f**), soma area (**g**) and TH expression (**h**). DA-Raptor KO;Rictor WT in blue: n=1999 neurons from four mice, DA-Raptor KO;Rictor KO in purple: n=1755 neurons from four mice, ****p<0.0001, Kolmogorov–Smirnov tests. **i,j)** Higher magnification merged images of the boxed VTA regions in panels b and c, showing TH (green) and p-S6 (magenta), scale bars = 50 μm. **k-m)** Cumulative distributions of VTA DA neuron p-S6 levels (**k**), soma area (**l**) and TH expression (**m**). DA-Raptor KO;Rictor WT in blue: n=2125 neurons from four mice, DA-Raptor KO;Rictor KO in purple: n=2976 neurons from four mice. ****p < 0.0001, Kolmogorov–Smirnov tests. **n)** Mean ± SEM [DA]_o_ versus time evoked from different striatal subregions elicited by a single electrical pulse. Traces are an average of n=24 (sites #1-6) or n=48 (site #7) transients per sampling region from six mice per genotype. DA-Raptor KO;Rictor WT in blue and DA-Raptor KO;Rictor KO in purple. Inset, typical cyclic voltammograms show characteristic DA waveform. **o)** Mean ± SEM peak [DA]_o_ by striatal subregion expressed as a percentage of DA-Raptor KO;Rictor WT (sampling region numbers correspond to the sites in **n**). n=24 (sites #1-6) or n=48 (site #7) transients per sampling region from six mice per genotype. p_1_ = 0.1208, p_3_ = 0.0691, p_5_ = 0.1875, *p_6_ = 0.0425, *p_7_ = 0.0002, Wilcoxon’s two-tailed t-tests; *p_2_ = 0.0238, *p_4_ = 0.0098, paired two-tailed t-tests. **p)** Mean ± SEM [DA]_o_ versus time averaged across all transients from three striatal territories, dorsal striatum (Str) (sites #1–3), central-ventral striatum (sites #4-6) and NAc core (site #7). Traces are an average of n=72 transients (sites #1-3, 4-6) or n=48 transients (site #7) per sampling territory from six mice per genotype. Statistical comparisons for the peak evoked [DA]_o_ between genotypes by sub-region: ***p_Str 1-3_=0.0004, **p_Str 4-6_=0.0019, ***p_NAc 7_ =0.0002, Wilcoxon’s two-tailed t-tests. **q)** Mean ± SEM [DA]_o_ versus time from concentration- and site-matched FCV transients recorded in dorsal and central-ventral striatum (sites #1-6). DA-Raptor KO;Rictor WT (“KO WT”) in blue and DA-Raptor KO;Rictor KO (“KO KO”) in purple. DA-Raptor KO;Rictor WT average of n=26 transients from six mice per genotype, DA-Raptor KO; Rictor KO average of n=24 transients from six mice per genotype. **r)** Single-phase exponential decay curve-fit of the falling phase of concentration- and site-matched striatal DA transients (from panel **q**). X-axis starts 375 ms after stimulation. DA-Raptor KO;Rictor WT average of n=26 transients from six mice per genotype, DA-Raptor KO;Rictor KO average of n=24 transients from six mice per genotype. ****p<0.0001, least-squares curve-fit comparison. **s)** Mean ± SEM [DA]_o_ versus time from concentration- and site-matched FCV transients recorded in NAc core (site #7). DA-Raptor KO;Rictor WT (“KO WT”) in blue and DA-Raptor KO;Rictor KO (“KO KO”) in purple. DA-Raptor KO;Rictor WT average of n=10 transients from five mice per genotype, DA-Raptor KO; Rictor KO average of n=9 transients from five mice per genotype. **t)** Single-phase exponential decay curve-fit of the falling phase of concentration-matched NAc DA transients (from panel **s**). X-axis starts 375 ms after stimulation onset. DA-Raptor KO;Rictor WT average of n=10 transients from five mice per genotype, DA-Raptor KO;Rictor KO average of n=9 transients from five mice per genotype. ****p<0.0001, least-squares curve-fit comparison.

To determine how DA neurotransmission was affected by combined suppression of mTORC1 and mTORC2 signaling, we measured evoked DA release with FCV. We found that compared to *Rptor* loss alone, deletion of *Rptor* and *Rictor* resulted in a further reduction in peak evoked DA levels (Figure 10n-p). Evoked DA release was reduced across all striatal sampling sites by ∼20% in Double KO mice compared to DA-Raptor KO mice (Figure 10o,p). Thus, concurrent mTORC1 and mTORC2 inhibition resulted in additional release deficits at dorsal striatum sites 1-3, which were unaffected by mTORC2 suppression alone (see Figure 7a-c). We examined the kinetics of DA re-uptake via DAT and found a significant reduction in the re-uptake rate in both the dorsal striatum (Figure 10q,r) and the NAc core (Figure 10s,t) of Double KO mice. In this case, combined mTORC1 and mTORC2 suppression caused slower [DA]_o_ uptake in the NAc core, which was not observed with either mTORC1 (see Figure 6f,g) or mTORC2 (see Figure 7f,g) inhibition alone. Together these data suggest that mTORC2 may have functions independent of mTORC1 that impact dopaminergic output. Combined inhibition of mTORC1 and mTORC2 signaling leads to widespread disruption of DA neuron morphology and output, highlighting the importance of mTOR signaling for dopamine neuron function.

## Discussion

DA modulates the activity of neural circuits throughout the brain and dysregulation of DA signaling is linked to a variety of neuropsychiatric and neurodegenerative disorders. The mTOR signaling network is a central regulator of multiple aspects of DA neuron growth and metabolism. Specifically, mTOR signaling has been shown to be important for therapeutic responses to PD, adaptations to drugs of abuse, and DA system changes induced by schizophrenia and ASD-linked mutations (Collo et al., 2013; Dadalko et al., 2015b; Kim et al., 2012; Kosillo and Bateup, 2021; Kosillo et al., 2019; Liu et al., 2018b; Malagelada et al., 2010; Mazei-Robison et al., 2011; Zhu et al., 2019). Investigating the DA neuron properties that are controlled by the mTOR pathway, and how these are regulated by the two mTOR complexes, is therefore critical for understanding how mTOR regulates DA neuron biology in both health and disease.

To address this, we genetically manipulated the activity of mTORC1 and mTORC2 in DA neurons and defined how these manipulations affect key cellular properties. Our results reveal several main findings. First, we find that chronic inhibition of mTORC1 signaling strongly impacts multiple aspects of DA neuron biology with broad effects on both SNc and VTA DA neurons. Second, we find that reducing mTORC2 signaling results in some phenotypes that overlap with mTORC1 inhibition. However, these phenotypes are generally milder and more selective. Third, mTORC2 inhibition leads to distinct cellular changes not observed following mTORC1 suppression, suggesting some independent actions of the two mTOR complexes in DA neurons. Fourth, we observe that concurrent disruption of both mTORC1 and mTORC2 signaling strongly compromises DA neuron output, underscoring the importance of mTOR for proper DA system function. Supplemental Table 3 presents a summary of the main findings of this study.

### mTORC1 is a potent regulator of DA neuron morphology and physiology

Here we found that chronic inhibition of mTORC1 signaling led to pronounced and widespread changes in the structure and function of midbrain DA neurons. Specifically, deletion of *Rptor* caused hypotrophy of multiple cellular compartments including the soma, dendrites and axons of both SNc and VTA neurons. Within the striatum we found that *Rptor* loss significantly reduced DA axon density and the size of individually resolved axons. Unexpectedly, we found that the striatal matrix compartment showed a greater DA axon innervation deficit compared to the patch regions in DA-Raptor KO mice. While selective degeneration of patch-projecting SNc DA neurons in PD models has been reported (Graybiel and Crittenden, 2011; Sgobio et al., 2017), we found that Raptor loss more strongly reduced DA projections to the striatal matrix. Two possible explanations for this relate to the developmental timing and trophic support of DA axons in patch versus matrix. Dopaminergic innervation of striatal patches is established first, forming so-called “DA islands” (Edley and Herkenham, 1984; Fishell and van der Kooy, 1987; Graybiel, 1984). It could be the case that matrix-projecting axonal segments, which develop later, fail to properly form in the context of limited molecular resources due to mTORC1 suppression. Further, compared to the matrix compartment, patches provide additional trophic support in the form of glia-derived neurotrophic factor (GDNF) (Lopez-Martin et al., 1999; Oo et al., 2005). This, in turn, could enable DA neurons with limited resources to better maintain axonal branches in striatal patches of DA-Raptor KO mice.

The somatodendritic hypotrophy of DA-Raptor KO neurons is consistent with a large body of literature demonstrating the importance of mTORC1 signaling in controlling cell size, including prior work showing that reduction of Raptor expression causes somatodendritic hypotrophy in other neuron types (Angliker et al., 2015; McCabe et al., 2020; Urbanska et al., 2012). While the mechanisms downstream of mTORC1 that regulate DA neuron size are not well understood, cellular hypotrophy of Raptor KO neurons may result from reduced protein and lipid synthesis, chronic activation of autophagy, and/or changes to the cytoskeleton (Fingar, 2002). A reduction in dendritic arborization would have implications for the number and/or type of synaptic inputs onto DA-Raptor KO neurons, which may impact their response to activation of upstream circuits. Consistent with their overall small size, DA-Raptor KO neurons exhibited changes in their passive membrane properties that rendered them hyperexcitable. Specifically, DA-Raptor KO neurons had increased membrane resistance, decreased rheobase, and increased action potential firing to small depolarizing currents. DA-Raptor KO neurons in the SNc also exhibited altered sag and rebound responses following a hyperpolarizing current injection. The latter changes could not be accounted for by altered passive properties, therefore they may have been driven by alterations in the expression of specific ion channels as seen in other cell types with mTOR manipulations (Raab-Graham et al., 2006).

Despite their increased intrinsic excitability, DA-Raptor KO neurons were severely impaired in their ability to release DA in response to electrical stimulation. This impairment was present across all dorsal and ventral striatal regions. In the dorsal striatum, which is primarily innervated by SNc DA neurons, we also observed a slower clearance rate of DA, which may be a compensatory change to counteract reduced DA neurotransmission. Compromised DA release in DA-Raptor KO mice was likely due to impaired DA synthesis since the expression of TH was reduced in striatal DA axons and in cell bodies. Correspondingly, we observed significantly reduced total tissue DA content in DA-Raptor KO mice. It is possible that mTORC1 is a direct regulator of TH protein synthesis as prior studies have shown that the translation of TH is controlled by cAMP (Chen et al., 2008) and that cAMP has a positive effect on translation via mTORC1 (Kim et al., 2010). This possibility is consistent with the bidirectional regulation of TH protein levels by mTORC1 signaling. Taken together, our data show that chronic developmental suppression of mTORC1 signaling in DA neurons leads to pronounced cellular hypotrophy, aberrant striatal innervation, and severe deficits in DA production and release.

### Chronic mTORC1 inhibition or activation impairs striatal DA release via different mechanisms

Previous studies have investigated how activation of mTORC1 signaling affects DA neuron structure and function. Chronic mTORC1 activation has been achieved via deletion of genes encoding upstream negative regulators such as Pten or Tsc1 (Diaz-Ruiz et al., 2009; Kosillo et al., 2019) or by constitutive activation of positive regulators such as Akt and Rheb (Cheng et al., 2011; Kim et al., 2012; Ries et al., 2006). In general, these studies find opposing phenotypes to what we observed following Raptor loss. Namely, mTORC1 activation leads to an increase in somatodendritic size and complexity, enlarged axon terminal area, enhanced axonal sprouting, and increased TH expression and DA synthesis (Cheng et al., 2011; Diaz-Ruiz et al., 2009; Kim et al., 2012; Kosillo et al., 2019; Ries et al., 2006). Nevertheless, with constitutive mTORC1 activation in Tsc1 KO neurons, evoked DA release in the dorsal striatum is reduced to a similar degree as observed with Raptor loss (Kosillo et al., 2019). Thus, while mTORC1 signaling bidirectionally controls DA neuron architecture and intrinsic excitability, either too much or too little mTORC1 activity is similarly detrimental to evoked dopaminergic neurotransmission.

In the case of Tsc1 loss, the evoked DA release deficits likely result from structural alterations in DA axon terminals that render vesicular DA release less efficient, since the total tissue DA content is elevated (Kosillo et al., 2019). In this context, the dorsal striatum is most strongly impacted as milder impairments are observed in evoked DA release in the ventral striatum, suggesting greater resilience of VTA DA axons to mTORC1 overactivation (Kosillo et al., 2019). By contrast, with Raptor loss, the DA release deficits are likely due to reduced TH levels and impaired DA synthesis, causing broad deficits across all DA neurons. Therefore, as observed in other neuronal types, both activation and inhibition of mTORC1 can produce similarly detrimental outcomes albeit by distinct mechanisms (Angliker et al., 2015).

### mTORC2 selectively alters the excitability and output of VTA DA neurons

The functions of mTORC2 have been less well studied; however, studies using mice with conditional deletion of *Rictor* from different neuron types have uncovered roles for mTORC2 in synaptic transmission, synaptic plasticity, actin dynamics and somatodendritic morphology (Angliker et al., 2015; Huang et al., 2013; McCabe et al., 2020; Thomanetz et al., 2013; Urbanska et al., 2012; Zhu et al., 2018). In terms of the DA system, conditional deletion of *Rictor* in VTA neurons using AAV-Cre has been shown to moderately decrease the soma size of DA neurons, similar to our observations (Mazei-Robison et al., 2011). It was further shown that chronic morphine treatment decreased VTA neuron soma size, increased spontaneous firing rate, but decreased electrically evoked DA release in the NAc (Mazei-Robison et al., 2011). The authors concluded that these cellular changes were driven by suppression of mTORC2 as they could be phenocopied by *Rictor* deletion and rescued by *Rictor* overexpression in the VTA (Mazei-Robison et al., 2011). This set of cellular changes, namely reduced soma size, increased excitability, and reduced dopaminergic output to the NAc is very similar to what we observed with DA neuron-specific deletion of *Rictor*. Here we also examined SNc DA neurons and found that like VTA neurons, they exhibit modest changes in somatodendritic architecture and reduced striatal axon density. However, in contrast to VTA neurons*, Rictor* KO SNc neurons did not show major changes in intrinsic excitability or DA release properties. Thus, VTA neurons are preferentially affected by suppression of mTORC2 signaling and SNc DA neurons are more resilient to mTORC2 perturbations.

It was previously reported that Nestin-Cre mediated deletion of *Rictor* from most neurons led to increased DAT expression and function together with reduced tissue DA content and amphetamine-stimulated release in the striatum, likely due to reduced striatal DA synthesis (Dadalko et al., 2015b). In the same mouse model, another study reported decreased DA content in the PFC, resulting from non-conventional DA uptake by norepinephrine transporter and its subsequent conversion to norepinephrine (Siuta et al., 2010). TH-Cre induced deletion of *Rictor* in catecholamine neurons was also shown to decrease DA tissue content the NAc (Dadalko et al., 2015a). Here we found that while DA neuron-specific deletion of *Rictor* reduced TH levels in the cell bodies of VTA DA neurons, this did not translate into significant changes in striatal TH expression or DA tissue content. This may reflect differences in the mouse models used as with Nestin- or TH-Cre driven deletion of *Rictor*, other neuron types may non-autonomously impact DA neurotransmission. Nonetheless, it is possible that the small but significant reduction in total DA axon density we observed, combined with reduced VTA TH expression and a trend toward lower striatal DA content could together compromise evoked DA release in DA-Rictor KO mice.

The morphological changes observed in DA-Rictor KO neurons at the level of cell bodies and axons were similar to DA-Raptor KO neurons, albeit smaller in magnitude. However, we found that loss of Rictor differentially affected the dendritic morphology of DA neurons. Specifically, while DA-Raptor KO neurons had reduced dendrite length and branching, Rictor KO DA neurons had increased dendritic complexity that was primarily driven by changes in proximal dendrites within 100 *μ*m of the soma. This increase in proximal dendrite branching is notably similar to the phenotype caused by Rictor loss from cerebellar Purkinje cells (PCs) (Thomanetz et al., 2013). In PCs, deletion of *Rictor* decreases soma size and axon diameter but increases the number of primary dendrites (Thomanetz et al., 2013). It was suggested that the effects of Rictor loss on dendritic morphology may be due to alterations in PKC levels or activity as PKC is a phosphorylation target of mTORC2 and a known regulator of actin cytoskeletal dynamics (Angliker and Ruegg, 2013). Further studies could investigate the potential role of PKC signaling in DA neurons in mediating the effects of Rictor loss on cytoskeletal organization.

### Crosstalk between mTORC1 and mTORC2 signaling

In this study, we used a genetic strategy to delete obligatory components of mTORC1 and mTORC2 that are specific for each complex. While deletion of *Rptor* or *Rictor* may initially disrupt the formation or activity of each complex selectively, we find that these manipulations do not exclusively affect the signaling of just one complex. Specifically, we find that deletion of *Rictor* from DA neurons leads to a small but significant reduction in p-S6 levels, a canonical mTORC1 read-out. While several studies have reported no change in mTORC1 phosphorylation targets with deletion or reduction of *Rictor* (Chen et al., 2019; Mazei-Robison et al., 2011; Thomanetz et al., 2013), other studies have shown reduced p-S6, similar to what we observe here (McCabe et al., 2020; Urbanska et al., 2012). This discrepancy could arise due to neuron type, developmental timing, and/or total duration and extent of Rictor loss.

In addition to changes in p-S6, we found a ∼10% reduction in soma size in DA-Rictor KO mice. Reduced soma size is a phenotype observed in multiple cell types with mTORC1 suppression. Therefore, it is possible that the reduction in soma size in Rictor KO DA neurons is due to the small reduction in mTORC1 activity, as determined by reduced p-S6 levels. Consistent with this idea, activation of Akt or p70S6K in cultured hippocampal neurons can restore p-S6 levels and rescue somatodendritic hypotrophy in cells treated with shRNA against *Rictor* (Urbanska et al., 2012). This suggests that some of the phenotypes associated with mTORC2 disruption may be driven by reduced mTORC1 activity. However, the reduced soma size of DA-Rictor KO neurons could arise from an mTORC1-independent mechanism. In support of the latter, it was shown that chronic morphine reduced the soma size of VTA DA neurons via an mTORC2-dependent but mTORC1-independent mechanism (Mazei-Robison et al., 2011). Such a conclusion is also supported by our observation that concurrent deletion of *Rptor* and *Rictor* from DA neurons led to a further reduction in soma size compared to loss of *Rptor* alone, suggesting independent mechanisms. We also observed a greater deficit in DA release in Raptor/Rictor Double KO mice compared to mice with loss of Raptor alone, further suggesting that mTORC1 and mTORC2 have independent functions in DA neurons. The mechanisms underlying the signaling interactions between mTORC1 and mTORC2 in DA neurons and other cell types are not well understood and warrant further investigation.

### ProExM-TrailMap enables quantitative measurement of DA axon density and size

A key challenge with measuring the anatomical properties of DA axons been difficulty in resolving the structure of individual DA axons within densely innervated regions, such as the striatum. In a classic study, single neuron reconstruction was performed to demonstrate the highly elaborate projection patterns of individual DA axon arbors within the striatum (Matsuda et al., 2009). Super resolution imaging approaches have also been applied to DA axons to address this challenge including 3D-SIM (Liu et al., 2018a). Crittenden and colleagues used expansion microscopy (ProExM) to resolve the unique architecture of dendrite-axon bundles in the substantia nigra called “striosome-dendron bouquets” (Crittenden et al., 2016). Here we extended this approach to the analysis of striatal DA axons by combining ProExM and volumetric light sheet imaging with the recently developed computational method TrailMap (Friedmann et al., 2020), which allows automatic identification and segmentation of axonal processes. With this pipeline we were able to measure the properties of individual DA axon segments within different striatal subregions and investigate how dopaminergic projection density and individual axon radius were altered by manipulations to mTORC1 and mTORC2 signaling.

We first validated this workflow in DA-Tsc1 KO mice, which have established DA axon terminal hypertrophy as measured previously by EM (Kosillo et al., 2019). Importantly, EM cannot readily assess axon density since measurements are made on cross-sections of individual axon terminals. In turn, quantifying bulk striatal TH immunofluorescence as a proxy of axon density is confounded by potential changes in TH expression, which is dynamically regulated (Chen et al., 2008). ProExM-TrailMap overcomes these challenges and enables accurate quantification of striatal DA axon volume to determine whether alterations in this measure are due to changes in axon density, size, or both.

Here, using ProExM-TrailMap, we find that mTORC1 manipulations bi-directionally control DA axon volume in the DLS and NAc core. Specifically, constitutive mTORC1 activation drives a substantial increase in DA axon density and radius, while chronic mTORC1 suppression significantly decreases axon density and radius. Notably, we observed regional heterogeneity of DA axons in DA-Raptor KO mice as putative patch regions exhibited increased axon density compared to the neighboring matrix, while axon radius was similarly reduced in both compartments. Constitutive mTORC2 inhibition selectively impacted DA axon density without affecting the radius of individual axon segments. These findings are important for understanding the impact of mTOR signaling manipulations on the DA axon network, which ultimately controls the functional output of the DA system.

### Summary and relevance to disorders affecting the DA system

Our findings reveal that mTORC1 manipulations have the most pronounced effects on DA neuron structure and function, while the consequences of mTORC2 disruption are more modest. Nonetheless, balanced mTOR signaling is critically important to dopaminergic output, which is compromised by either chronic up- or down-regulation of mTORC1 or suppression of mTORC2 signaling. An important consideration is that the developmental timing and total duration of mTOR signaling manipulation will likely influence how changes in mTOR affect DA output. For example, altering mTOR signaling in embryonic development, as done in this study, likely leads to reduced DA output early in development, enabling post-synaptic circuits to compensate for low dopaminergic tone. This may allow for normal motor behavior in the face of significant deficits in DA output (Delignat-Lavaud et al., 2021; Kosillo et al., 2019). By contrast, it is known that abruptly reducing DA output in adulthood leads to significant motor impairments as observed in PD (Iancu et al., 2005). In addition, short-versus long-term mTOR signaling manipulations may lead to distinct outcomes. For example, in PD, acute upregulation of mTOR signaling via constitutively active Akt or Rheb may be beneficial to cell survival and axonal growth but prolonged activation, as with embryonic *Tsc1* or *Pten* deletion, is detrimental to DA output (Diaz-Ruiz et al., 2009; Domanskyi et al., 2011; Kosillo et al., 2019; Zhu et al., 2019). Similarly, acute (partial) inhibition of mTOR with rapamycin can suppress synthesis of pro-apoptotic proteins and may promote removal of misfolded proteins and damaged organelles by increasing autophagy (Hernandez et al., 2012; Malagelada et al., 2010). This may improve cell survival in response to neurotoxic injury. However, our results show that chronic mTORC1 suppression is detrimental to the health and output of DA neurons. These factors will need to be taken into account when considering mTOR modulators as potential therapies.

In summary, this work, together with prior studies, demonstrates the importance of mTORC1 and mTORC2 signaling for proper DA neuron morphology, physiology and output and reveals how unbalanced mTOR signaling can profoundly affect DA neuron structure and function.

## Methods

### Mice

All animal procedures and husbandry were carried out in accordance with protocols approved by the University of California, Berkeley Institutional Animal Care and Use Committee (IACUC). Both male and female animals were used for all experiments. In general, young adult mice between 2-4 months of age were used, the ages of the animals used are indicated in the methods for each experiment. Mice were housed with same sex littermates in groups of 5-6 animals per cage and kept on a regular 12 hr light/dark cycle (lights on at 7am), with ad libitum access to food and water.

To generate DA neuron-specific Raptor-KO mice, *Rptor^fl/fl^* mice (Jackson Laboratories strain #013188) were bred to *DAT-Cre^wt/+^*mice (Jackson Laboratories strain *#*0066600), on a C56Bl/6J background. A *Rptor^wt/fl^;DAT-Cre^wt/+^* × *Rptor^wt/fl^;DAT-Cre^wt/+^* cross was used to generate experimental animals. Experimental mice were heterozygous for *DAT-Cre* and either homozygous WT (*Rptor^wt/wt^;DAT-Cre^wt/+^*, referred to as DA-Raptor WT) or homozygous floxed for *Rptor* (*Rptor^fl/fl^;DAT-Cre^wt/+^*, referred to as DA-Raptor KO).

To generate DA neuron-specific Rictor-KO mice, *Rictor^fl/fl^* mice (Jackson Laboratories strain #020649) were bred to *DAT-Cre^wt/+^* mice (Jackson Laboratories strain *#*0066600), on a mixed genetic background (C57Bl/ 6J, 129 SvJae and BALB/cJ). A *Rictor^wt/fl^; DAT-Cre^wt/+^* × *Rictor^wt/fl^; DAT-Cre^wt/+^*cross was used to generate experimental animals. Experimental mice were heterozygous for *DAT-Cre* and either homozygous WT (*Rictor^wt/wt^;DAT-Cre^wt/+^*, referred to as DA-Rictor WT) or homozygous floxed for *Rictor* (*Rictor^fl/fl^;DAT-Cre^wt/+^*, referred to as DA-Rictor KO).

To generate DA neuron-specific double KO mice, *Rptor^fl/fl^* mice were crossed to *Rictor^fl/fl^* mice and their offspring were bred to *DAT-Cre^wt/+^* mice. These mice were on mixed genetic background (C57Bl/ 6J, CD1, 129 SvJae and BALB/cJ). To generate experimental animals a *Rptor^fl/fl^;Rictor^wt/fl^;DAT-Cre^wt/wt^* × *Rptor^fl/fl^;Rictor^wt/fl^;DAT-Cre^+/+^* cross was used. Experimental mice were heterozygous for *DAT-Cre*, homozygous for floxed *Rptor,* and either homozygous WT for *Rictor* (*Rptor^fl/fl^;Rictor^wt/wt^;DAT-Cre^wt/+^*, referred to as DA-Raptor KO;Rictor WT) or homozygous floxed for *Rictor* (*Rptor^fl/fl^;Rictor^fl/fl^;DAT-Cre^wt/+^*, referred to as DA-Raptor KO;Rictor KO).

For cell counting and axonal arborization experiments, *Rptor^fl/fl^;DAT-Cre^wt/+^* and *Rictor^fl/fl^;DAT-Cre^wt/+^* mice were bred to the Ai9 tdTomato Cre-reporter line (Jackson Laboratories strain #007909). Mice used for cell counting experiments were either heterozygous or homozygous for the Ai9 transgene. For ProExM analysis, only heterozygous Ai9 animals were used.

### Neurobiotin-filled neuron reconstruction

Male and female mice (P56-P80) were deeply anesthetized by isoflurane, transcardially perfused with ice cold high Mg^2+^ ACSF using a perilstatic pump (Instech) and decapitated. 275 *μ*m thick coronal midbrain slices were prepared on a vibratome (Leica VT1000 S) in ice cold high Mg^2+^ ACSF containing in mM: 85 NaCl, 25 NaHCO_3_, 2.5 KCl, 1.25 NaH_2_PO_4_, 0.5 CaCl_2_, 7 MgCl_2_, 10 glucose, and 65 sucrose. Slices recovered for 15 minutes at 34°C followed by at least 50 minutes at room temperature (RT) in ACSF containing in mM: 130 NaCl, 25 NaHCO_3_, 2.5 KCl, 1.25 NaH_2_PO_4_, 2 CaCl_2_, 2 MgCl_2_, and 10 glucose. All solutions were continuously bubbled with 95% O_2_ and 5% CO_2_. For whole cell recordings, 2.5-6 mΩ borosilicate glass pipettes (Sutter Instrument: BF150-86-7.5) were filled with a potassium-based internal solution containing in mM: 135 KMeSO_3_, 5 KCl, 5 HEPES, 4 Mg-ATP, 0.3 Na-GTP, 10 phospho-creatine, 1 EGTA, and 4mg/ml neurobiotin (Vector laboratories #SP-1120).

275 μm slices containing dopamine neurons filled with neurobiotin-containing internal solution (4 mg/ml) during patch-clamp experiments were fixed in 4% paraformaldehyde solution (Electron Microscopy Sciences: 15713) in 1x PBS for 24-48 h at 4°C. With continuous gentle shaking, slices were washed in 1x PBS 5 x 5 min and incubated with BlockAid blocking solution (Life Tech: B10710) for 1 h at RT. Primary tyrosine hydroxylase antibody (1:1000, Immunostar: 22941) and streptavidin Alexa Flour 488 conjugate (Invitrogen: S11223) were applied overnight at 4°C in 1x PBS containing 0.25% (v/v) Trinton-X-100 (PBS-Tx). The following day, slices were washed in 1x PBS 5 x 5 min, and Alexa-633 goat anti-mouse secondary antibody (1:500, ThermoFisher: A21050) in PBS-Tx was applied for 1 h at RT. Slices were washed in cold 1x PBS 5 x 5 min, mounted on SuperFrost slides (VWR: 48311-703) with the cell-containing side of the slice facing up, and coverslipped with either Prolong Gold antifade (Life Tech: P36935) or Vectashield hard-set (Vector Labs: H-1500) mounting media.

Filled cells in the mounted sections were imaged on a Zeiss LSM 880 NLO AxioExaminer confocal with 20x/1.0 N.A. water immersion objective and 488 nm Argon laser using 1.53 µm steps to acquire a z-stack image spanning the entirety of the neurobiotin-filled cell body and dendritic arbor. 3D reconstruction of the cells was done using IMARIS 9.2.1 software (Bitplane) with automated filament tracing and manual editing. The mask generated by the automated filament tracing algorithm was continuously cross-referenced with the original z-stack image to ensure accuracy. Spurious segments created by the automated filament tracer due to background noise were removed, while processes with incomplete reconstruction were edited to incorporate missing segments.

### Immunohistochemistry

Male and female mice (P75-P90) were deeply anesthetized by isoflurane and transcardially perfused with ice cold 1x PBS (∼5-7 ml) followed by 4% paraformaldehyde (PFA) solution (Electron Microscopy Sciences: 15713) in 1x PBS (∼5-10 ml) using a perilstatic pump (Instech). The brains were removed and post-fixed by immersion in 4% PFA in 1x PBS solution overnight at 4°C. Brains were suspended in 30% sucrose in 0.1 M PB for cryoprotection. After brains descended to the bottom of the vial (typically 24-28 h), 30 μm coronal sections of the midbrain and striatum were cut on a freezing microtome (American Optical AO 860), collected into serial wells, and stored at 4°C in 1x PBS containing 0.02% (w/v) sodium azide (NaN_3_; Sigma Aldrich). All animals used for histology experiments were heterozygous for the Ai9 tdTomato Cre-reporter allele.

Brain sections for immunohistochemistry were batch processed to include matched control and experimental animals from a given line. Free-floating sections were washed with gentle shaking, 3 x 5 min in 1x PBS followed by 1 h incubation at RT with BlockAid blocking solution (Life Tech: B10710). Primary antibodies were applied at 4°C in 1x PBS containing 0.25% (v/v) Trinton-X-100 (PBS-Tx) for 48-72 h to ensure penetration of the antibody throughout the slice. Sections were then washed with cold 1x PBS 3 x 5 min and incubated for 1 h at RT with secondary antibodies in PBS-Tx. Sections were washed in cold 1x PBS 5 x 5 min, mounted on SuperFrost slides (VWR: 48311-703), and coverslipped with Vectashield hard-set (Vector Labs: H-1500) mounting media.

The following primary antibodies were used: tyrosine hydroxylase (1:1000, Immunostar: 22941) and phospho-S6 (Ser240/244) (1:1000, Cell Signaling: 5364S). The following secondary antibodies were used: Alexa-488 goat anti-mouse (1:1000, Thermo Fisher: A-11001) and Alexa-633 goat anti-rabbit (1:1000, Thermo Fisher: A-11034).

### Confocal microscopy

Images of 30 μm sections processed for immunohistochemistry were acquired using a Zeiss LSM 880 NLO AxioExaminer confocal microscope fitted with a motorized XY-stage for tile scanning. A 20x/1.0 N.A. water objective was used to generate tile scans (scan area 425.1 x 425.1 μm per tile) of the entire midbrain (9 x 3 grid). 488 nm, 561 nm and 633 nm lasers were used. Z-stack images captured the entire thickness of the slice at 1.70 μm steps. Laser power and gain settings were kept constant between control and experimental slice imaging.

### Electrophysiology

Male and female adult mice (P56–P80) were deeply anesthetized by isoflurane, transcardially perfused with ice-cold high Mg^2+^ artificial cerebrospinal fluid (aCSF) using a peristaltic pump (Instech) and decapitated. Coronal midbrain slices (275 μm thick) were prepared on a vibratome (Leica VT1000 S) in ice-cold high Mg^2+^ aCSF containing (in mM): 85 NaCl, 25 NaHCO3, 2.5 KCl, 1.25 NaH2PO4, 0.5 CaCl2, 7 MgCl2, 10 glucose, and 65 sucrose. Slices were recovered for 15 min at 34°C, followed by at least 50 min at room temperature (RT) in aCSF containing (in mM): 130 NaCl, 25 NaHCO3, 2.5 KCl, 1.25 NaH2PO4, 2 CaCl2, 2 MgCl2, and 10 glucose. All solutions were continuously bubbled with 95% O_2_ and 5% CO_2_.

Recordings were performed at 32°C in the presence of AMPA, NMDA, and GABA-A receptor blockers (10 μM GYKI 52466 cat #1454, 10 μM CPP cat #0247, 50 μM picrotoxin cat #1128, final concentrations, all from Tocris), with a bath perfusion rate of ∼2 ml/min. Dopaminergic neurons in the SNc and VTA were identified by tdTomato fluorescence. For whole-cell recordings, 2.5–6 mΩ borosilicate glass pipettes (Sutter Instruments: BF150-86-7.5) were filled with a potassium-based internal solution containing (in mM): 135 KMeSO3, 5 KCl, 5 HEPES, 4 Mg-ATP, 0.3 Na-GTP, 10 phospho-creatine, 1 EGTA, and 4 mg/ml neurobiotin (Vector Laboratories #SP-1120). Recordings were obtained using a MultiClamp 700B amplifier (Molecular Devices) and ScanImage software (https://github.com/bernardosabatini/SabalabAcq). Passive membrane properties were recorded in voltage clamp with the membrane held at −70 mV. Negative current steps (2 s, −50 to −200 pA) were applied to measure the sag amplitude and rebound following hyperpolarization. Positive current steps (2 s, +25 to +600 pA) were applied to generate an input–output curve from a baseline membrane potential of −70 mV as maintained by current clamp. For whole-cell recordings, series resistance was <30 MΩ and liquid junction potential was not corrected.

Electrophysiology data were acquired using the ScanImage software, written and maintained by Dr. Bernardo Sabatini (https://github.com/bernardosabatini/SabalabAcq). Data were analyzed in Igor Pro (Wavemetrics). Rheobase was calculated as the current at which APs were first elicited. Passive properties were calculated from an RC check in voltage clamp recordings at −70 mV.

### Fast-scan cyclic voltammetry (FCV)

DA release was monitored using fast-scan cyclic voltammetry (FCV) in acute coronal slices. Male and female mice (P56-P90) were deeply anesthetized by isoflurane and decapitated. 275 *μ*m thick coronal striatal slices were prepared on a vibratome (Leica VT1000 S) in ice cold high Mg^2+^ ACSF containing in mM: 85 NaCl, 25 NaHCO_3_, 2.5 KCl, 1.25 NaH_2_PO_4_, 0.5 CaCl_2_, 7 MgCl_2_, 10 glucose, and 65 sucrose. Slices recovered for 1 h at RT and were recorded from in ACSF containing in mM: 130 NaCl, 25 NaHCO_3_, 2.5 KCl, 1.25 NaH_2_PO_4_, 2 CaCl_2_, 2 MgCl_2_, and 10 glucose. All solutions were continuously saturated with 95% O_2_ and 5% CO_2._ Slices between +1.5 mm and +0.5 mm from bregma containing dorsal striatum and nucleus accumbens were used for experimentation (Franklin and Paxinos, 2008). Slices from both genotypes were prepared and recorded from on the same day, the order of brain dissection was counterbalanced between experiments.

In the recording chamber, slices were maintained at 32°C with a superfusion rate of 1.2-1.4 ml/min. Slices used for recording were anatomically matched between animals and location within the recording chamber was counterbalanced between experiments. Extracellular DA concentration ([DA]_o_) evoked by local electrical stimulation was monitored with FCV at carbon-fiber microelectrodes (CFMs) using a Millar voltammeter (Julian Millar, Barts and the London School of Medicine and Dentistry). CFMs were fabricated in-house from epoxy-free carbon fiber ∼7 μm in diameter (Goodfellow Cambridge Ltd) encased in a glass capillary (Harvard Apparatus: GC200F-10) pulled to form a seal with the fiber and cut to final tip length of 70–120 μm. The CFM was positioned ∼ 100 μm below the tissue surface at a 45° angle. A triangular waveform was applied to the carbon fiber scanning from −0.7 V to +1.3 V and back, against an Ag/AgCl reference electrode at a rate of 800 V/s. Evoked DA transients were sampled at 8 Hz, and data were acquired at 50 kHz using AxoScope 10.5-10.7 (Molecular Devices). Oxidation currents evoked by electrical stimulation were converted to [DA]_o_ from post-experimental calibrations. Recorded FCV signals were confirmed to be DA by comparing oxidation (+0.6 V) and reduction (−0.2 V) potential peaks from experimental voltammograms with currents recorded during calibration with 2 μM DA dissolved in ACSF.

A concentric bipolar stimulating electrode (FHC: CBAEC75) used for electrical stimulation was positioned on the slice surface with minimal tissue disturbance within 100 μm of the CFM. Following 30 min slice equilibration in the recording chamber, DA release was evoked using square wave pulses (0.6 mA pulse amplitude, 2 ms pulse duration) controlled by an Isoflex stimulus isolator (A.M.P.I., Jerusalem, Israel) delivered out of phase with voltammetric scans. At each sampling site, DA release was evoked every 2.5 min. Three stimulations were delivered (in the following order: single pulse, 4 pulses at 100 Hz, single pulse) before progressing to the corresponding site in the other slice.

### High-performance liquid chromatography (HPLC)

Tissue DA content was measured by HPLC with electrochemical detection in tissue punches from dorsal and ventral striatum. Male and female mice (P60-P90) were deeply anesthetized by isoflurane and decapitated. 300 *μ*m thick coronal slices of striatum were prepared on a vibratome (Leica VT1000 S) in ice cold high Mg^2+^ ACSF containing in mM: 85 NaCl, 25 NaHCO_3_, 2.5 KCl, 1.25 NaH_2_PO_4_, 0.5 CaCl_2_, 7 MgCl_2_, 10 glucose, and 65 sucrose. Slices recovered for 1 hour at RT in ACSF containing in mM: 130 NaCl, 25 NaHCO_3_, 2.5 KCl, 1.25 NaH_2_PO_4_, 2 CaCl_2_, 2 MgCl_2_, and 10 glucose. All solutions were continuously bubbled with 95% O_2_ and 5% CO_2_. Following slice recovery, tissue punches from the dorsal (2.5 mm diameter) and ventral striatum/NAc (1.5 mm diameter) from two brain slices per animal were taken and stored at -80°C in 200 µl 0.1 M HClO_4_. On the day of analysis, samples were thawed, homogenized by sonication, and centrifuged at 16,000 x g for 15 min at 4°C. The supernatant was analyzed for DA content using HPLC with electrochemical detection. Analytes were separated using a 4.6 x 150 mm Microsorb C18 reverse-phase column (Varian or Agilent) and detected using a Decade II SDS electrochemical detector with a Glassy carbon working electrode (Antec Leyden) set at +0.7 V with respect to a Ag/AgCl reference electrode. The mobile phase consisted of 13% methanol (v/v), 0.12 M NaH2PO4, 0.5 mM OSA, 0.8 mM EDTA, pH 4.8, and the flow rate was fixed at 1 ml/min. Analyte measurements were normalized to tissue punch volume (pmol/mm^3^). HPLC analysis was repeated in two independent experiments. Raw HPLC measurements were normalized to WT control values for each line to enable percent-change quantification of DA and DOPAC availability in KO samples. Raw values are reported in Supplemental Table 2.

### Western blotting (WB)

Male and female mice (P60-P90) were deeply anesthetized by isoflurane and decapitated. Bilateral striata were rapidly dissected out on ice, flash-frozen in liquid nitrogen and stored at - 80°C. On the day of analysis, frozen samples were sonicated until homogenized (QSonica Q55) in 500 *μ*l lysis buffer containing 1% SDS in 1x PBS with Halt phosphatase inhibitor cocktail (Fisher: PI78420) and Complete mini EDTA-free protease inhibitor cocktail (Roche: 4693159001). Sample homogenates were then boiled on a heat block at 95°C for 10 min, allowed to cool to RT and total protein content was determined by BCA assay (Fisher: PI23227). Following the BCA assay, protein homogenates were mixed with 4x Laemmli sample buffer (Bio-Rad: 161-0747) and 10-15 μg of protein were loaded onto 4–15% Criterion TGX gels (Bio-Rad: 5671084) in running buffer (3.03 g Tris base, 14.41 g glycine, 1 g SDS in 1 L ultrapure dH_2_O). Proteins were transferred to a PVDF membrane (BioRad: 1620177) in transfer buffer (3.03 g Tris base, 14.41 g glycine in 1 L ultrapure dH_2_O) at 4°C overnight using a Bio-Rad Criterion Blotter (12 V constant voltage). The membranes were blocked in 5% milk in 1x TBS with 1% Tween (TBS-T) for one hour at RT, and incubated with primary antibodies diluted in 5% milk in TBS-T overnight at 4°C. The following day, after 3 x 10 min washes with TBS-T, the membranes were incubated with HRP-conjugated secondary antibodies (1:5000 dilution in 5% milk in TBS-T solution) for 1h at RT. Following 6 x 10 min washes, the membranes were incubated with chemiluminesence substrate (Perkin-Elmer: NEL105001EA) for 1 min and signal was developed on GE Amersham Hyperfilm ECL (VWR: 95017-661). Membranes were stripped by 2 x 7 min incubations in stripping buffer (6 M guanidine hydrochloride (Sigma: G3272) with 1:150 β-mercaptoethanol) with shaking, followed by 4 x 2 min washes in 1x TBS with 0.05% NP-40 to re-blot on subsequent days.

The following primary antibodies were used: mouse anti-Tyrosine Hydroxylase (1:2000, Immunostar: 22941); rabbit anti-DARPP32 (1:1500, Cell Signaling: 2306S); mouse anti-Histone- 3 (1:1500, Cell Signaling: 96C10); rabbit anti-VMAT2 (1:1000, Alomone Labs: AMT-006); mouse anti-DAT (1:1000, Abcam: 128848) and rabbit anti DAT phospho-T53 (1:1000, Abcam: 183486). Secondary antibodies were goat anti-rabbit HRP (1:5000, Bio-Rad: 170-5046) and goat anti-mouse HRP (1:5000, Bio-Rad: 170-5047). Bands were quantified by densitometry using FIJI (NIH). Phospho-proteins were normalized to their respective total proteins. Total proteins were normalized to Histone 3 loading control.

### Protein-retention expansion microscopy (ProExM)

PFA fixed coronal 30 μm striatal brain sections were washed 3 x 5 min in 1x PBS and followed by 1 h incubation with Blockaid at RT with gentle nutation. Rabbit anti-RFP antibody (1:500; Rockland: 600-401-379) was applied at 4°C in 1x PBS containing 0.25% Triton-X-100 (PBS-Tx) for 24 h. Sections were washed with 1x PBS 5 x 5 min, incubated for 24 h at 4°C with a goat anti-rabbit Alexa 546 secondary antibody (1:500; Thermo: A-11035) in PBS-Tx, and washed with cold PBS 5 x 5 min.

Sections were then processed with ProExM protocol as previously described (Asano et al., 2018; Tillberg et al., 2016). Briefly, sections were incubated with anchoring solution for 16 h at RT with no shaking. The anchoring solution consisted of 10 mg/mL acryloyl-X SE (Thermo-Fisher: A20770) dissolved in anhydrous DMSO (Thermo-Fisher: D12345) diluted to 0.1 mg/mL in 1x PBS. Slices were then washed 2 x 15 min in 1x PBS. Monomer solution consisting of 1x PBS, 2 M NaCl, 8.55% (w/v) sodium acrylate (Combi-Blocks: QC-1489), 2.5% (w/v) acrylamide (Sigma: A9099), and 0.15% (w/v) N,N′-methylenebisacrylamide (Invitrogen: M7279) was frozen in aliquots and warmed to 4°C before use. A gelling solution was made by combining monomer solution with 0.01% (w/v) 4-hydroxy-TEMPO (4HT, Sigma 176141), 0.2% (w/v) tetra-methylethylenediamine (TEMED, Sigma: T7024) and 0.2% (w/v) ammonium persulfate (APS, Sigma: A3678). All chemical concentrations refer to their final ratio in the gelling solution (monomer solution with 4HT, TEMED, and APS).

Slices were incubated in the gelling solution for 30 min at 4°C, transferred to gelation chambers (22 x 25 x 0.15 mm) and placed in a humidity chamber for 2 h at 37°C for polymerization (see Figure 4 in (Asano et al., 2018) for depiction of the chamber). Gels were then placed in Proteinase K (New England Biolabs: P8107S) diluted 1:100 (8 U/mL) in digestion buffer composed of 50 mM Tris (pH 8), 1 mM EDTA, 0.5% (w/v) Triton X-100, 1 M NaCl for 16 h at RT and then stored in 1x PBS at 4°C.

Post-expansion imaging of DA axons was carried out as follows. No. 1.5 coverslip strips cut to 2.75 x 0.5 mm with a diamond scriber (VWR: 52865-005) were cleaned with distilled water followed by pure ethanol before 20 min treatment with 0.1% (w/v) poly-L-lysine. Following 3 rinses with distilled water, coverslips were left to air-dry for 1 hour. Coverslips were superglued onto the inset surface of a custom-made light-sheet adapter (Asano et al., 2018) printed on a Form 2 (Formlabs) using standard resin. Expansion gels were incubated 6 x 15 min in distilled water, and following full expansion were trimmed to obtain regions containing dorsolateral striatum or ventromedial striatum and nucleus accumbens core (regions of interest, ROI). Trimmed gels containing ROIs were carefully placed onto the poly-L-lysine-coated coverslip strip and set aside to adhere for 5 min. Gels were imaged on a Zeiss Lightsheet Z.1 using a 20x/1.0 N.A. water immersion objective and a single-side illumination 10x/0.2 N.A. objective. Images were acquired with a 546 nm laser line using 0.522 µm steps to acquire a z-stack image of entire expanded region of interest.

To quantify the expansion factor, pre- and post-expansion images of the same field of view were acquired on an Olympus FV3000 confocal with a 4x/0.16 N.A. air objective. Expansion factor calculation was performed using an implementation of the scale-invariant feature transform (SIFT) algorithm (Lowe, 2004). OpenCV package for Python (https://github.com/opencv/opencv-python) was used to generate SIFT keypoints and estimate a partial 2D affine transformation between pre- and post-expansion keypoints, restricting image alignment to rotation, translation, and uniform scaling. Analysis code is available at https://github.com/kamodulin/expansion-microscopy.

### TrailMap axon segmentation

To identify DA axons within striatal sub-regions, expansion microscopy volumes were segmented with a convolutional neural network, TrailMap (Friedmann et al., 2020), with a few modifications. Three separate volumes were acquired for each ROI, two of which were held out for inference (428 volumes total). The remaining volumes were randomly assigned to training (88 volumes) or validation (25 volumes) datasets and the rest were not used. 3D sub-volumes (400 x 400 pixels) that spanned the entire depth of the original volume were randomly cropped from training and validation datasets. These volumes were sparsely annotated with a spacing of ∼30-50 slices along the Z axis. From these sub-volumes, 64^3^ pixel cubes were randomly cropped to generate 8,800 training and 2,500 validation volumes. The model was trained with real-time data augmentation and the model with the lowest validation loss (epoch 22) was selected for inference on the main dataset. Our best trained model and a fork of the original TrailMap package with modified data augmentation procedure are available at https://github.com/kamodulin/TRAILMAP.

TrailMap predictions were thresholded at *P* > 0.7 to generate binary volumes and objects less than 256 voxels in size were removed. Total axon volume was calculated relative to the total image volume. Binary volumes were skeletonized in three dimensions and the total length of the axon armature was measured and normalized by the total image volume. A 3D Euclidean distance transform was computed on the binary volume for every point along the axon skeleton. The average of these distances was used to calculate the mean axon radius per image. All measurements were normalized by expansion factor. Analysis code is available at https://github.com/kamodulin/expansion-microscopy.

### Striatal patch-matrix ROI processing

PFA fixed coronal 30 μm striatal brain sections were washed 3 x 5 min in 1x PBS and followed by 1 h incubation with Blockaid (Life Tech - B10710) at RT with gentle rocking. Rabbit anti-mu opioid receptor (MOR) antibody (1:1000; EMD Millipore: AB5511) was applied at 4°C in 1x PBS containing 0.25% Triton-X-100 (PBS-Tx) for 24 h. Sections were washed with 1x PBS 5 x 5 min, incubated for 1 h at RT with a goat anti-rabbit Alexa 488 secondary antibody (Thermo: A-110) in PBS-Tx, and washed with cold 1x PBS 5 x 5 min. Processed sections were mounted on SuperFrost slides (VWR: 48311-703), and coverslipped with Prolong Gold antifade (Life Tech: P36935) mounting media.

Striatal sections containing tdTomato labelled DA axons and with patch (striosome) regions visualized by high density MOR signal were imaged on an Olympus FV3000 confocal microscope equipped with 488 and 633 nm lasers and a motorized stage for tile imaging. Z stack images captured the entire thickness of the section at 2.75 μm steps with a 10x air/0.4 N.A. objective (UPLXAPO10X).

Patch ROIs were manually drawn around the high intensity MOR regions in Image J (NIH). The same ROIs were then moved outside the high intensity MOR signal to get fluorescence measurements in striatal matrix. Quantification of tdTomato fluorescence intensity signal for patch and matrix ROIs was done on max-projected Z stack images.

### Quantification and statistical analyses

Whenever possible, quantification and analyses were performed blind to genotype. Statistical analyses and graphing were performed using the GraphPad Prism 8 software. All datasets were first analyzed using D’Agostino and Pearson normality test, and then parametric or non-parametric two-tailed statistical tests were employed accordingly to determine significance. If the variances between two groups was significantly different, a Welch’s correction was applied. Significance was set as *p < 0.05, **p < 0.01, ***p < 0.001, and ****p < 0.0001. P values were corrected for multiple comparisons. Statistical details for each experiment are reported in the figure legends.

## Acknowledgements

This work was supported by NIH grant #R01NS105634 to H.S.B. P.K. was supported by a post-doctoral fellowship from the Tuberous Sclerosis Alliance (#381490). H.S.B. and P.K. were supported by NARSAD Young Investigator Grants from the Brain and Behavior Research Foundation (#25073 to H.S.B. and #27458 to P.K.). H.S.B. is a Chan Zuckerberg Biohub Investigator. B.M.R. was supported by Clarendon Fund Studentship. This work was also supported by Parkinson’s UK (Grant G-1504). Confocal and light sheet imaging experiments were conducted at the CRL Molecular Imaging Center, RRID:SCR_017852, supported by the Helen Wills Neuroscience Institute. We would like to thank Holly Aaron and Feather Ives for their microscopy advice and support. We thank Mary (Tien) Chiu and Victoria Du for their assistance in maintaining the mouse colony. We thank Dr. Drew Friedmann for his advice in adapting the TrailMap method to our analysis pipeline. We thank the members of the Bateup lab for their feedback on this work.

## Conflicts of interest

The authors declare no conflicts of interest.

---

**Supplemental Figure 1.**
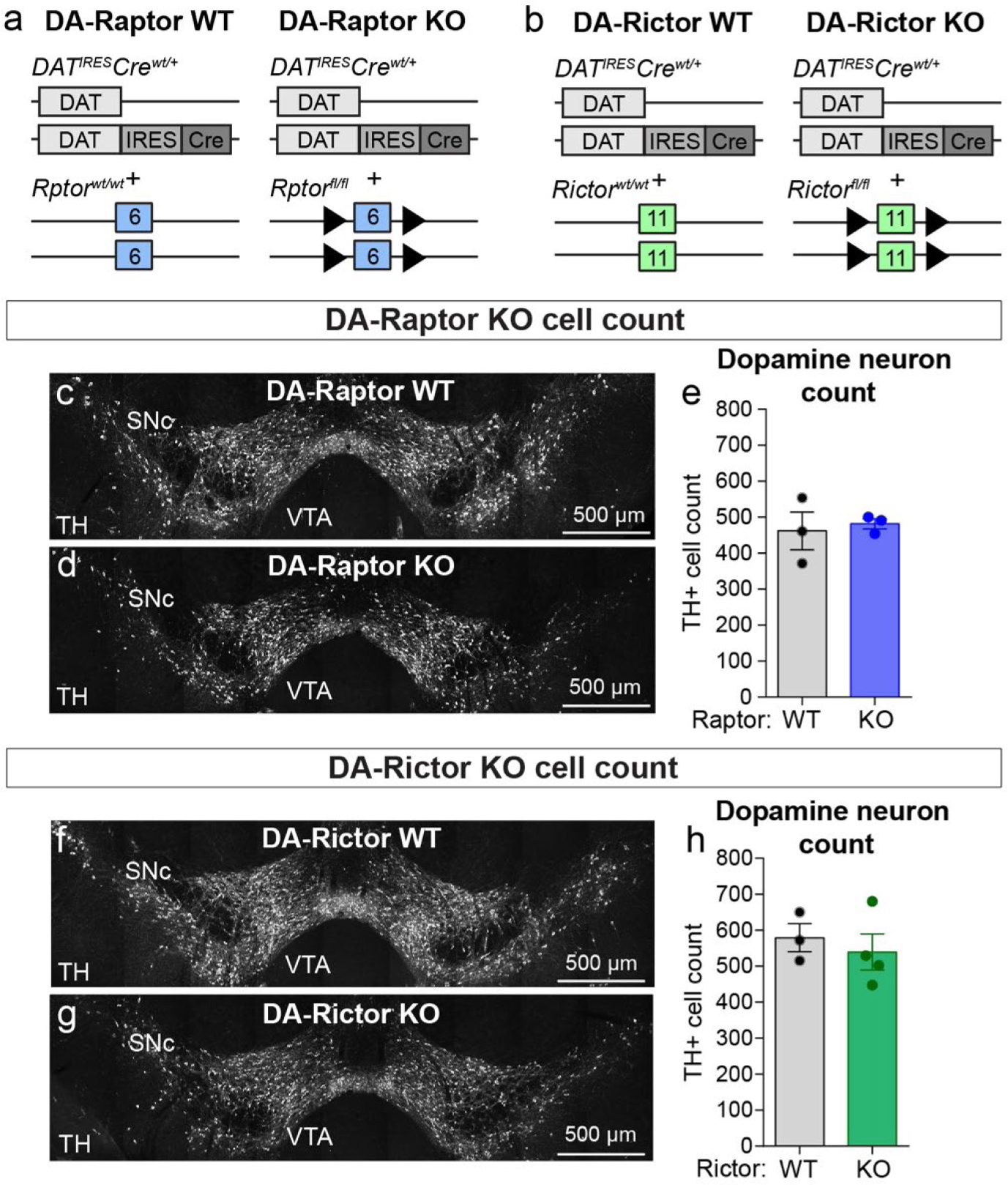
Deletion of *Rptor* or *Rictor* does not affect DA neuron number (related to Figures 1 and 2). **a,b)** Schematic of the genetic strategies to delete *Rptor* (**a**) or *Rictor* (**b**) selectively from DA neurons. Numbered boxes represent exons and triangles represent loxP sites. **c,d)** Representative confocal images of midbrain sections from DA-Raptor WT (**c**) and DA-Raptor KO (**d**) mice with DA neurons visualized by tyrosine hydroxylase (TH) immunostaining, scale bars=500 μm. **e)** Mean ± SEM number of TH-positive DA neurons in the midbrain (SNc and VTA combined) in DA-Raptor WT and KO mice. n=3 mice per genotype (averaged from two sections per mouse), p=0.7484, Welch’s two-tailed t-test. **f,g)** Representative confocal images of midbrain sections from DA-Rictor WT (**f**) and DA-Rictor KO (**g**) mice with DA neurons visualized by TH immunostaining, scale bars=500 μm. **h)** Mean ± SEM number of TH-positive DA neurons in the midbrain (SNc and VTA combined) in DA-Rictor WT and KO mice. n=3 DA-Rictor WT and 4 DA-Rictor KO mice (averaged from two sections per mouse), p=0.5587, Welch’s two-tailed t-test. For all bar graphs, dots represent values for individual mice.

**Supplemental Figure 2.**
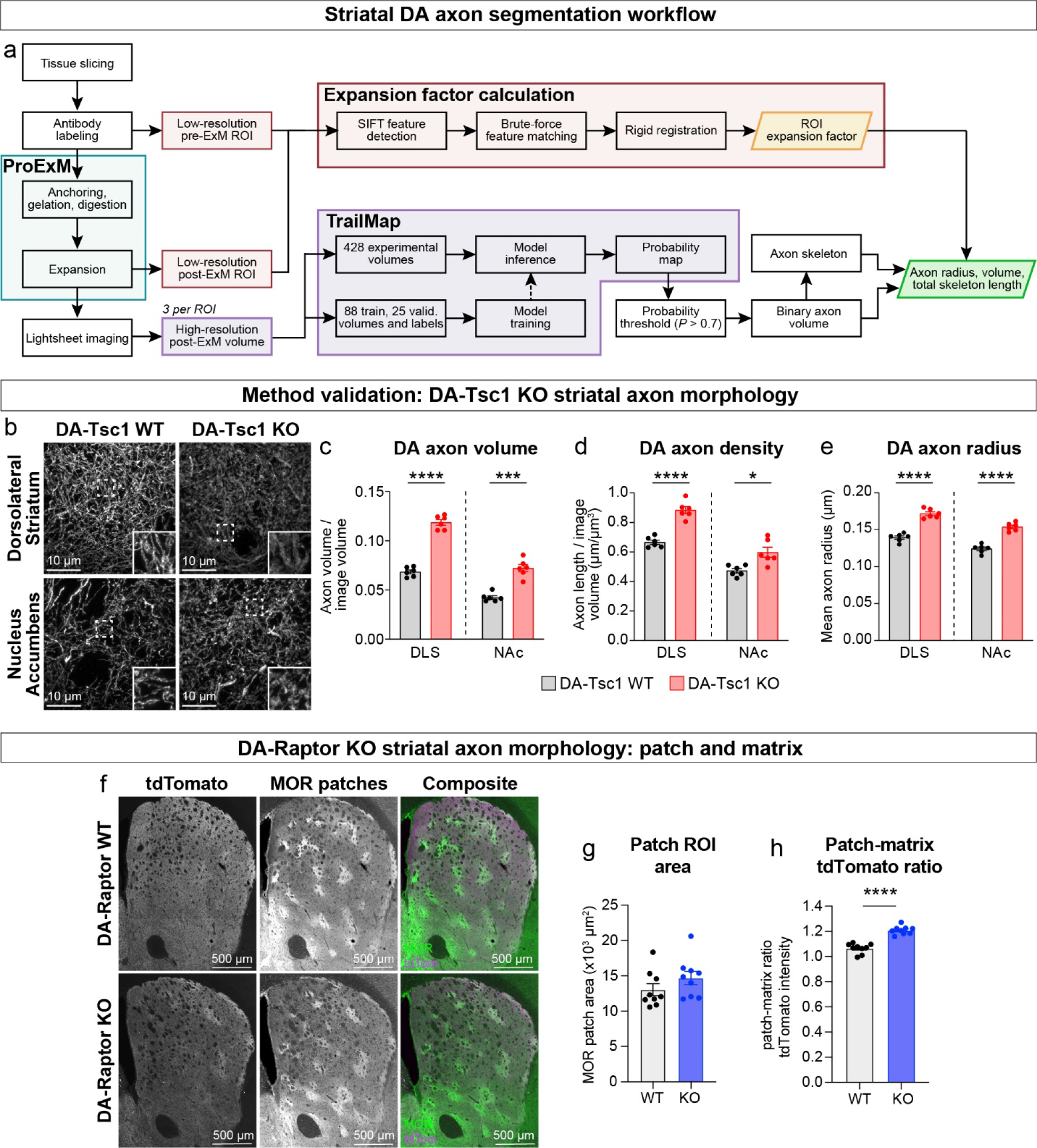
Development of a ProExM-TrailMap pipeline for the analysis of DA axon morphology (related to Figure 3). **a)** Workflow for performing protein-retention expansion microscopy (ProExM), light sheet imaging and TrailMap segmentation and analysis of striatal DA axons. **b)** Representative light sheet images of striatal DA axons visualized by Cre-dependent tdTomato fluorescence enhanced with anti-RFP antibody. Images show ProExM tissue gels expanded by a factor of ∼4.5. Scale bars are normalized by expansion factor. Insets are higher magnification images of boxed regions from the same panels. **c-e)** Mean ± SEM DA axon volume (**c**), density (**d**) and radius (**e**) in the dorsolateral striatum (DLS) and nucleus accumbens core (NAc) regions from DA-Tsc1 WT and DA-Tsc1 KO mice. n=6 slices from 3 mice per genotype (values are the average of 4 images per slice). Axon volume (**c**), ****p<0.0001 WT vs KO in DLS, ***p=0.0001 WT vs KO in NAc. Axon density (**d**), ****p<0.0001 WT vs KO in DLS, *p=0.0123 WT vs KO in NAc. Axon radius (**e**), ****p<0.0001 WT vs KO in DLS, ****p<0.0001 WT vs KO in NAc. Welch’s two-tailed t-tests. **f)** Representative confocal images of striatal sections from DA-Raptor WT (top panels) and DA-Raptor KO (bottom panels) mice with striatal DA axons visualized by Cre-dependent tdTomato and striatal patch (striosome) regions labeled with an antibody against μ-opioid receptor (MOR), scale bars=500 μm. Composite images show TH in green and tdTomato in magenta. **g,h)** Mean ± SEM total patch area (**g**) and patch/matrix ratio of tdTomato fluorescence intensity (**h**) in the striatum of DA-Raptor WT and DA-Raptor KO mice. n=9 sections from three mice per genotype. Patch area (**g**), p=0.2054, Welch’s two-tailed t-test. Patch/matrix ratio (**h**), ****p<0.0001, Welch’s two-tailed t-test. For all bar graphs, dots represent individual striatal sections. See also Supplemental Videos 1-7

**Supplemental Figure 3.**
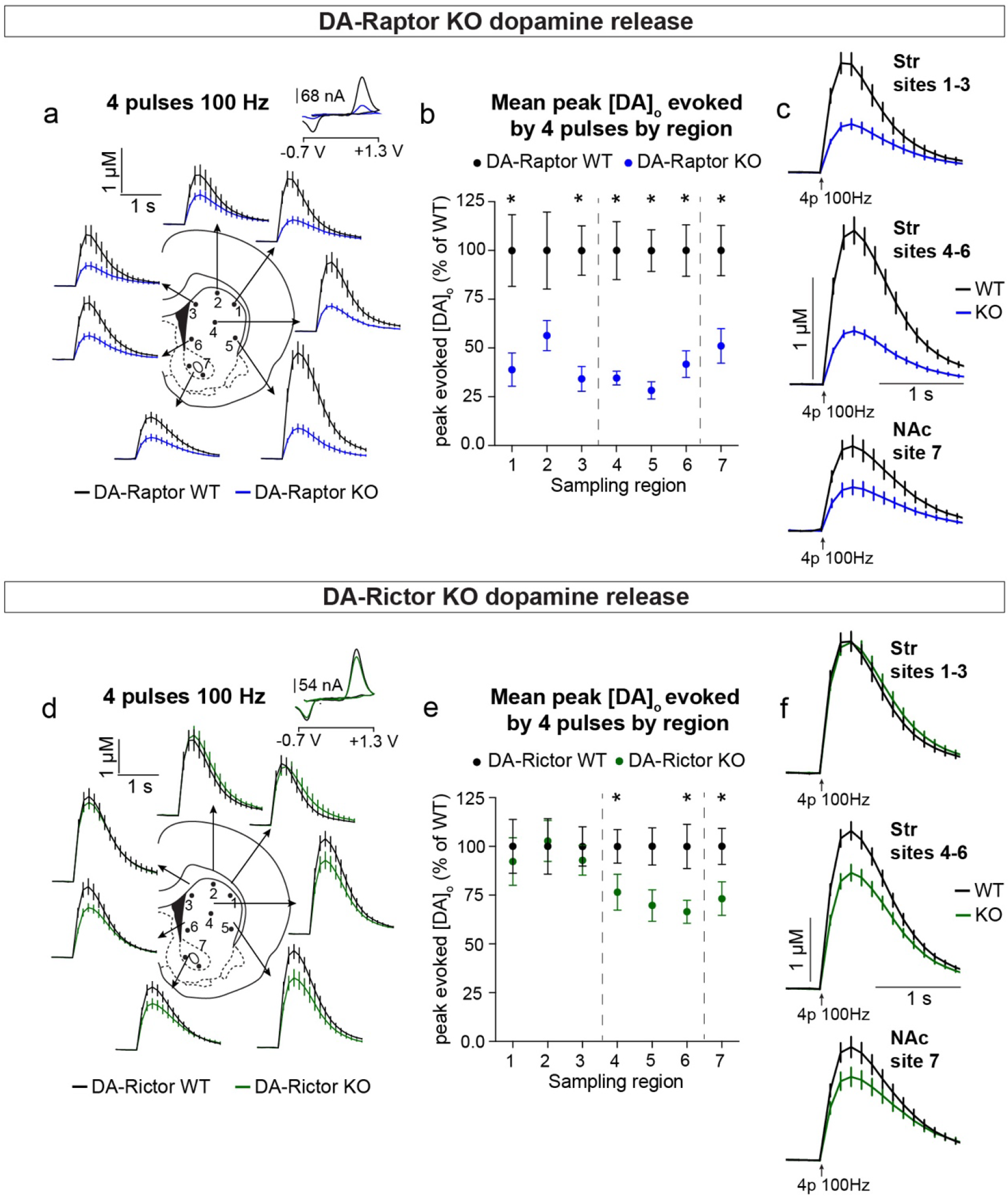
mTORC1 and mTORC2 inhibition differentially impact evoked striatal DA release (related to Figures 6 and 7). **a)** Mean ± SEM [DA]_o_ versus time evoked from different striatal subregions by a high frequency train of 4 pulses delivered at 100 Hz. Traces are an average of n=10 (sites #1,2,4), n=9 (sites #3,6), n=8 (site #5) and n=17 transients (site #7) per sampling region from five mice per genotype. DA-Raptor WT in black, DA-Raptor KO in blue. Inset, typical cyclic voltammograms show characteristic DA waveform. **b)** Mean ± SEM peak [DA]_o_ by striatal subregion expressed as a percentage of WT (sampling region numbers correspond to the sites in panel a). n=10 (sites #1,2,4), n=9 (sites #3,6), n=8 (site #5) and n=17 transients (site #7) per sampling region from five mice per genotype. *p_1_ = 0.0273, *p_3_ = 0.0078, *p_4_ = 0.0020, Wilcoxon’s two-tailed t-tests; *p_2_ = 0.0826; *p_5_ = 0.0004, *p_6_ = 0.0021, *p_7_ = 0.0022, paired two-tailed t-tests. **c)** Mean ± SEM [DA]_o_ versus time averaged across all transients from three striatal territories, dorsal striatum (Str) (sites #1-3), central-ventral striatum (sites #4-6) and NAc core (site #7). Traces are an average of n=29 (sites #1-3), n=27 (sites #4-6) and n=17 transients (site #7) per sampling territory from five mice per genotype. Statistical comparisons for the peak evoked [DA]_o_ between genotypes by subregion: ****p_Str 1-3_<0.0001, ****p_Str 4-6_<0.0001, ****p_NAc 7_=0.0022, paired two-tailed t-tests. **d)** Mean ± SEM [DA]_o_ versus time evoked from different striatal subregions by a high frequency train of 4 pulses delivered at 100 Hz. Traces are an average of n=10 (sites #1-6) or n=20 (site #7) transients per sampling region from five mice per genotype. DA-Rictor WT in black, DA-Rictor KO in green. Inset, typical cyclic voltammograms show characteristic DA waveform. **e)** Mean ± SEM peak [DA]_o_ by striatal subregion expressed as a percentage of WT (sampling region numbers correspond to the sites in panel d). n=10 (sites #1-6) or n=20 (site #7) transients per sampling region from five mice per genotype. p_1_ = 0.4354, p_2_ = 0.8331, p_3_ = 0.4252, *p_4_ = 0.0281; p_5_ = 0.0548, *p_6_ = 0.0042, *p_7_ = 0.0031, paired two-tailed t-tests. **f)** Mean ± SEM [DA]_o_ versus time averaged across all transients from three striatal territories, dorsal striatum (Str) (sites #1-3), central-ventral striatum (sites #4-6) and NAc core (site #7). Traces are an average of n=30 (sites #1-3, 4-6) and n=20 transients (site #7) per sampling territory from five mice per genotype. Statistical comparisons for the peak evoked [DA]_o_ between genotypes by subregion: p_Str 1-3_=0.4522, Wilcoxon’s two-tailed t-test; ****p_Str 4-6_<0.0001, **p_NAc 7_ =0.0031, paired two-tailed t-tests.

## Supplemental Video Titles and Legends

**Supplemental Video 1. 3D rendering of tdTomato-labeled DA axons in the dorsolateral striatum (related to Figures 3 and S2).**

Image shows a 3D rendering of tdTomato-labeled Raptor KO DA axons (*Rptor^fl/fl^;DAT-Cre^wt/+^;Ai9^wt/+^*) in the dorsolateral striatum (DLS). Tissue sections were expanded ∼4x using ProExM. Video shows a 41.87 x 41.87 x 50.52 μm volume generated from z-stack lightsheet images (normalized by expansion factor).

**Supplemental Video 2. Video of z-stack lightsheet images of tdTomato-labeled Tsc1 WT and Tsc1 KO DA axons in the DLS (related to Figures 3 and S2).**

Striatal sections containing tdTomato-labeled DA axons from DA-Tsc1 WT (*Tsc1^wt/wt^;DAT-Cre^wt/+^;Ai9^wt/+^*) and DA-Tsc1 KO (*Tsc1^fl/fl^;DAT-Cre^wt/+^;Ai9^wt/+^*) mice were expanded with ProExM and imaged on a lightsheet microscope. Video shows 38.64 x 38.64 x 32.07 μm (DA-Tsc1 WT) and 41.25 x 41.25 x 34.25 μm (DA-Tsc1 KO) z-stacks (normalized by expansion factor) from the DLS.

**Supplemental Video 3. Video of z-stack lightsheet images of tdTomato-labeled Tsc1 WT and Tsc1 KO DA axons in the nucleus accumbens core (related to Figures 3 and S2).**

Striatal sections containing tdTomato-labeled DA axons from DA-Tsc1 WT (*Tsc1^wt/wt^;DAT- Cre^wt/+^;Ai9^wt/+^*) and DA-Tsc1 KO (*Tsc1^fl/fl^;DAT-Cre^wt/+^;Ai9^wt/+^*) mice were expanded with ProExM and imaged on a lightsheet microscope. Video shows 39.32 x 39.32 x 25.13 μm (DA-Tsc1 WT) and 40.84 x 40.84 x 26.10 μm (DA-Tsc1 KO) z-stacks (normalized by expansion factor) from the nucleus accumbens core (NAc).

**Supplemental Video 4. Video of z-stack lightsheet images of tdTomato-labeled Raptor WT and Raptor KO DA axons in the DLS (related to Figures 3 and S2).**

Striatal sections containing tdTomato-labeled DA axons from DA-Raptor WT (*Rptor^wt/wt^;DAT-Cre^wt/+^;Ai9^wt/+^*) and DA-Raptor KO (*Rptor^fl/fl^;DAT-Cre^wt/+^;Ai9^wt/+^*) mice were expanded with ProExM and imaged on a lightsheet microscope. Video shows 40.97 x 40.97 x 35.60 μm (DA-Raptor WT) and 41.87 x 41.87 x 36.39 μm (DA-Raptor KO) z-stacks (normalized by expansion factor) from the DLS. For DA-Raptor KO sections, putative patch and matrix regions were imaged separately.

**Supplemental Video 5. Video of z-stack lightsheet images of tdTomato-labeled Raptor WT and Raptor KO DA axons in the NAc (related to Figures 3 and S2).**

Striatal sections containing tdTomato-labeled DA axons from DA-Raptor WT (*Rptor^wt/wt^;DAT-Cre^wt/+^;Ai9^wt/+^*) and DA-Raptor KO (*Rptor^fl/fl^;DAT-Cre^wt/+^;Ai9^wt/+^*) mice were expanded with ProExM and imaged on a lightsheet microscope. Video shows 41.65 x 41.65 x 35.45 μm (DA-Raptor WT) and 42.08 x 42.08 x 35.81 μm (DA-Raptor KO) z-stacks (normalized by expansion factor) from the NAc.

**Supplemental Video 6. Video of z-stack lightsheet images of tdTomato-labeled Rictor WT and Rictor KO DA axons in the DLS (related to Figures 3 and S2).**

Striatal sections containing tdTomato-labeled DA axons from DA-Rictor WT (*Rictor^wt/wt^;DAT-Cre^wt/+^;Ai9^wt/+^*) and DA-Rictor KO (*Rictor^fl/fl^;DAT-Cre^wt/+^;Ai9^wt/+^*) mice were expanded with ProExM and imaged on a lightsheet microscope. Video shows 39.62 x 39.62 x 30.17 μm (DA-Rictor WT) and 44.05 x 44.05 x 33.54 μm (DA-Rictor KO) z-stacks (normalized by expansion factor) from the DLS.

**Supplemental Video 7. Video of z-stack lightsheet images of tdTomato-labeled Rictor WT and Rictor KO DA axons in the NAc (related to Figures 3 and S2).**

Striatal sections containing tdTomato-labeled DA axons from DA-Rictor WT (*Rictor^wt/wt^;DAT-Cre^wt/+^;Ai9^wt/+^*) and DA-Rictor KO (*Rictor^fl/fl^;DAT-Cre^wt/+^;Ai9^wt/+^*) mice were expanded with ProExM and imaged on a lightsheet microscope. Video shows 39.19 x 39.19 x 36.52 μm (DA-Rictor WT) and 42.58 x 42.58 x 39.67 μm (DA-Rictor KO) z-stacks (normalized by expansion factor) from the NAc.

## Notes

### Competing Interest Statement

The authors have declared no competing interest.

