## Supplemental Table 1 for "Dopamine neuron morphology and output are differentially controlled by mTORC1 and mTORC2"

**Supplemental Table 1. Summary of the electrophysiology properties of SNc and VTA DA neurons, related to Figures 4 and 5.**

| | **DA-Raptor WT (SNc)** | | | | **DA-Raptor KO (SNc)** | | | | **WT vs KO** |
|---|---|---|---|---|---|---|---|---|---|
| properties | Mean | SEM | n (cells) | n (mice) | Mean | SEM | n (cells) | n (mice) | p-value/ test |
| **Series resistance** (mOhms) | 3.269 | 0.308 | 28 | 6 | 3.494 | 0.213 | 28 | 6 | 0.0965 Mann-Whitney |
| **Membrane resistance** (mOhms) | 85.79 | 7.869 | 28 | 6 | 321.0 | 22.20 | 28 | 6 | **<0.0001** Mann-Whitney |
| **Membrane capacitance** (pF) | 115.8 | 5.365 | 28 | 6 | 48.92 | 1.661 | 28 | 6 | **<0.0001** unpaired t-test |
| **Resting membrane potential** (mV) | -56.13 | 0.904 | 28 | 6 | -53.29 | 1.372 | 28 | 6 | 0.0902 unpaired t-test |
| **Rheobase** (current when first action potentials occur, pA) | 156.3 | 21.54 | 28 | 6 | 66.67 | 8.333 | 27 | 6 | **<0.0001** Mann-Whitney |
| **Action potential (AP) threshold** (mV) | -33.45 | 0.976 | 28 | 6 | -32.56 | 1.313 | 27 | 6 | 0.5885 unpaired t-test |
| **Action potential peak** (maximum membrane potential, mV) | 24.85 | 1.279 | 28 | 6 | 15.55 | 1.478 | 27 | 6 | **<0.0001** unpaired t-test |
| **Action potential height** (change in membrane potential from the start of the AP to maximum depolarization, mV) | 66.44 | 1.556 | 28 | 6 | 57.74 | 1.468 | 27 | 6 | **0.0002** unpaired t-test |
| **Afterhyperpolarization** (minimum membrane potential after the AP, mV) | -61.70 | 1.645 | 28 | 6 | -56.17 | 1.365 | 27 | 6 | **0.0127** unpaired t-test |
| **Afterhyperpolarization** (change in membrane potential from the start of the AP to maximum hyperpolarization, mV) | 20.11 | 1.171 | 28 | 6 | 15.06 | 1.011 | 27 | 6 | **0.0020** unpaired t-test |
| **Maximum hyperpolarization in response to -100 pA** (from ~-70 mV in response to a 2 second -100 pA current step, mV) | -91.04 | 1.841 | 28 | 6 | -124.4 | 2.271 | 28 | 6 | **<0.0001** unpaired t-test |
| **Sag component in response to -100 pA** (maximum hyperpolarization minus the steady state membrane potential in | 11.49 | 1.060 | 28 | 6 | 19.38 | 1.283 | 28 | 6 | **<0.0001** unpaired t-test |

| | | | | | | | | | |
|---|---|---|---|---|---|---|---|---|---|
| the last 50 ms of the current step, mV) | | | | | | | | | |
| **Sag component expressed as a percentage** (sag component as a percentage of the total step size, calculated as the difference between the max hyperpolarization and baseline potential, %) | 48.99 | 2.219 | 28 | 6 | 36.18 | 2.472 | 28 | 6 | **0.0003** Mann-Whitney |
| **Rebound depolarization in response to -100 pA** (baseline membrane potential minus the maximum depolarization within 500 ms of the end of the current step, mV) | 11.32 | 1.350 | 28 | 6 | 10.15 | 1.030 | 28 | 6 | 0.4946 unpaired t-test |
| **Rebound depolarization expressed as a percentage** (rebound as a percentage of the total step size, calculated as the difference between the max hyperpolarization and baseline potential, %) | 50.77 | 6.156 | 28 | 6 | 19.28 | 1.961 | 28 | 6 | **<0.0001** Mann-Whitney |
| | **DA-Raptor WT (VTA)** | | | | **DA-Raptor KO (VTA)** | | | | **WT vs KO** |
| properties | Mean | SEM | n (cells) | n (mice) | Mean | SEM | n (cells) | n (mice) | p-value/ test |
| **Series resistance** (mOhms) | 4.216 | 0.282 | 22 | 8 | 4.472 | 0.337 | 22 | 6 | 0.8617 Mann-Whitney |
| **Membrane resistance** (mOhms) | 463.7 | 34.41 | 22 | 8 | 485.9 | 40.15 | 22 | 6 | 0.6769 unpaired t-test |
| **Membrane capacitance** (pF) | 64.88 | 5.117 | 22 | 8 | 43.21 | 2.581 | 22 | 6 | **0.0005** unpaired t-test |
| **Resting membrane potential** (mV) | -52.03 | 2.130 | 22 | 8 | -51.42 | 3.046 | 22 | 6 | 0.8697 unpaired t-test |
| **Rheobase** (current when first action potentials occur, pA) | 69.32 | 6.565 | 22 | 8 | 63.64 | 16.75 | 22 | 6 | **0.0190** Mann-Whitney |
| **Action potential threshold** (mV) | -26.06 | 1.458 | 22 | 8 | -32.35 | 1.391 | 22 | 6 | **0.0039** Mann-Whitney |

| | | | | | | | | |
|---|---|---|---|---|---|---|---|---|
| **Action potential peak** (maximum membrane potential, mV) | 17.65 | 2.180 | 22 | 8 | 10.50 | 1.708 | 22 | 6 | **0.0134** unpaired t-test |
| **Action potential height** (change in membrane potential from the start of the AP to maximum depolarization, mV) | 54.72 | 2.378 | 22 | 8 | 53.49 | 1.735 | 22 | 6 | 0.6766 unpaired t-test |
| **Afterhyperpolarization** (minimum membrane potential after the AP, mV) | -51.09 | 1.619 | 22 | 8 | -54.24 | 0.875 | 22 | 6 | 0.0942 unpaired t-test |
| **Afterhyperpolarization** (change in membrane potential from the start of the AP to maximum hyperpolarization, mV) | 14.02 | 0.826 | 22 | 8 | 11.83 | 0.703 | 22 | 6 | 0.0983 Mann-Whitney |
| **Maximum hyperpolarization in response to -100 pA** (from ~-70 mV in response to a 2 second -100 pA current step, mV) | -128.9 | 4.016 | 22 | 8 | -148.4 | 4.986 | 21 | 6 | **0.0038** unpaired t-test |
| **Sag component in response to -100 pA** (maximum hyperpolarization minus the steady state membrane potential in the last 50 ms of the current step, mV) | 14.34 | 3.756 | 22 | 8 | 15.78 | 3.329 | 21 | 6 | 0.9329 Mann-Whitney |
| **Sag component expressed as a percentage** (sag component as a percentage of the total step size, calculated as the difference between the max hyperpolarization and baseline potential, %) | 20.33 | 3.465 | 22 | 8 | 19.18 | 3.388 | 21 | 6 | 0.3660 Mann-Whitney |
| **Rebound depolarization in response to -100 pA** (baseline membrane potential minus the maximum depolarization within 500 ms of the end of the current step, mV) | 2.414 | 0.595 | 22 | 8 | 2.027 | 0.753 | 21 | 6 | 0.6219 Mann-Whitney |
| **Rebound depolarization** | 3.857 | 0.777 | 22 | 8 | 2.944 | 1.177 | 21 | 6 | 0.2513 |

| | | | | | | | | | |
|---|---|---|---|---|---|---|---|---|---|
| **expressed as a percentage** (rebound as a percentage of the total step size, calculated as the difference between the max hyperpolarization and baseline potential, %) | | | | | | | | | Mann-Whitney |
| | **DA-Rictor WT (SNc)** | | | | **DA-Rictor KO (SNc)** | | | | **WT vs KO** |
| properties | Mean | SEM | n (cells) | n (mice) | Mean | SEM | n (cells) | n (mice) | p-value/ test |
| **Series resistance** (mOhms) | 2.979 | 0.249 | 21 | 4 | 2.573 | 0.171 | 27 | 8 | 0.1249 Mann-Whitney |
| **Membrane resistance** (mOhms) | 110.9 | 14.05 | 21 | 4 | 147.9 | 14.52 | 27 | 8 | 0.0772 Mann-Whitney |
| **Membrane capacitance** (pF) | 114.4 | 6.016 | 21 | 4 | 85.41 | 5.165 | 27 | 8 | **0.0006** Mann-Whitney |
| **Resting membrane potential** (mV) | -55.46 | 1.337 | 21 | 4 | -52.27 | 1.565 | 27 | 8 | 0.1419 unpaired t-test |
| **Rheobase** (current when first action potentials occur, pA) | 101.2 | 11.11 | 21 | 4 | 112.0 | 12.96 | 27 | 8 | 0.7414 Mann-Whitney |
| **Action potential threshold** (mV) | -32.94 | 1.520 | 21 | 4 | -31.34 | 0.974 | 27 | 8 | 0.9180 Mann-Whitney |
| **Action potential peak** (maximum membrane potential, mV) | 25.46 | 1.647 | 21 | 4 | 23.61 | 1.429 | 27 | 8 | 0.3964 unpaired t-test |
| **Action potential height** (change in membrane potential from the start of the AP to maximum depolarization, mV) | 66.31 | 1.253 | 21 | 4 | 62.61 | 1.616 | 27 | 8 | 0.1590 Mann-Whitney |
| **Afterhyperpolarization** (minimum membrane potential after the AP, mV) | -59.49 | 2.276 | 21 | 4 | -58.46 | 1.416 | 27 | 8 | 0.5467 Mann-Whitney |
| **Afterhyperpolarization** (change in membrane potential from the start of the AP to maximum hyperpolarization, mV) | 18.65 | 1.517 | 21 | 4 | 19.45 | 1.063 | 27 | 8 | 0.6582 unpaired t-test |
| **Maximum hyperpolarization in response to -100 pA** (from ~-70 mV in response to a 2 second -100 pA current step, mV) | -94.91 | 2.162 | 21 | 4 | -101.0 | 2.079 | 27 | 8 | **0.0490** unpaired t-test |

| | | | | | | | | | |
|---|---|---|---|---|---|---|---|---|---|
| **Sag component in response to -100 pA** (maximum hyperpolarization minus the steady state membrane potential in the last 50 ms of the current step, mV) | 11.90 | 0.992 | 21 | 4 | 14.74 | 1.129 | 27 | 8 | 0.0772 Mann-Whitney |
| **Sag component expressed as a percentage** (sag component as a percentage of the total step size, calculated as the difference between the max hyperpolarization and baseline potential, %) | 45.96 | 2.701 | 21 | 4 | 43.63 | 2.172 | 27 | 8 | 0.5006 unpaired t-test |
| **Rebound depolarization in response to -100 pA** (baseline membrane potential minus the maximum depolarization within 500 ms of the end of the current step, mV) | 14.17 | 4.150 | 21 | 4 | 12.02 | 1.363 | 27 | 8 | 0.8049 Mann-Whitney |
| **Rebound depolarization expressed as a percentage** (rebound as a percentage of the total step size, calculated as the difference between the max hyperpolarization and baseline potential, %) | 52.27 | 12.87 | 21 | 4 | 37.70 | 4.586 | 27 | 8 | 0.4212 Mann-Whitney |
| | **DA-Rictor WT (VTA)** | | | | **DA-Rictor KO (VTA)** | | | | **WT vs KO** |
| properties | Mean | SEM | n (cells) | n (mice) | Mean | SEM | n (cells) | n (mice) | p-value/ test |
| **Series resistance** (mOhms) | 3.608 | 0.303 | 22 | 4 | 4.239 | 0.446 | 20 | 6 | 0.2418 unpaired t-test |
| **Membrane resistance** (mOhms) | 485.9 | 46.92 | 22 | 4 | 606.2 | 43.56 | 20 | 6 | **0.0418** Mann-Whitney |
| **Membrane capacitance** (pF) | 60.06 | 3.977 | 22 | 4 | 47.28 | 4.014 | 20 | 6 | **0.0014** Mann-Whitney |

| | | | | | | | | | |
|---|---|---|---|---|---|---|---|---|---|
| **Resting membrane potential** (mV) | -49.72 | 1.700 | 22 | 4 | -50.45 | 2.267 | 20 | 6 | 0.9305 Mann-Whitney |
| **Rheobase** (current when first action potentials occur, pA) | 100.0 | 20.76 | 20 | 4 | 57.50 | 8.331 | 20 | 6 | **0.0299** Mann-Whitney |
| **Action potential threshold** (mV) | -26.61 | 2.422 | 20 | 4 | -24.32 | 1.568 | 19 | 6 | 0.4376 unpaired t-test |
| **Action potential peak** (maximum membrane potential, mV) | 19.25 | 2.135 | 20 | 4 | 14.25 | 2.431 | 19 | 6 | 0.1298 unpaired t-test |
| **Action potential height** (change in membrane potential from the start of the AP to maximum depolarization, mV) | 60.49 | 2.102 | 20 | 4 | 54.17 | 2.303 | 19 | 6 | **0.0496** unpaired t-test |
| **Afterhyperpolarization** (minimum membrane potential after the AP, mV) | -54.66 | 1.367 | 20 | 4 | -52.49 | 0.858 | 19 | 6 | 0.1935 unpaired t-test |
| **Afterhyperpolarization** (change in membrane potential from the start of the AP to maximum hyperpolarization, mV) | 14.15 | 1.135 | 20 | 4 | 12.57 | 0.883 | 19 | 6 | 0.2829 unpaired t-test |
| **Maximum hyperpolarization in response to -100 pA** (from ~-70 mV in response to a 2 second -100 pA current step, mV) | -135.2 | 5.016 | 22 | 4 | -143.5 | 4.862 | 20 | 6 | 0.2433 unpaired t-test |
| **Sag component in response to -100 pA** (maximum hyperpolarization minus the steady state membrane potential in the last 50 ms of the current step, mV) | 14.20 | 1.589 | 22 | 4 | 16.73 | 3.457 | 20 | 6 | 0.7178 Mann-Whitney |
| **Sag component expressed as a percentage** (sag component as a percentage of the total step size, calculated as the difference between the max hyperpolarization and baseline potential, %) | 22.13 | 2.189 | 22 | 4 | 20.35 | 3.093 | 20 | 6 | 0.3888 Mann-Whitney |
| **Rebound depolarization in response to -100 pA** (baseline membrane potential minus the | 3.42 | 0.573 | 22 | 4 | 2.720 | 0.661 | 20 | 6 | 0.4252 unpaired t-test |

| | | | | | | | | | |
|---|---|---|---|---|---|---|---|---|---|
| maximum depolarization within 500 ms of the end of the current step, mV) | | | | | | | | | |
| **Rebound depolarization expressed as a percentage** (rebound as a percentage of the total step size, calculated as the difference between the max hyperpolarization and baseline potential, %) | 6.044 | 1.290 | 22 | 4 | 3.515 | 0.808 | 20 | 6 | 0.1638 Mann-Whitney |
