## Supplemental Table 2 for "Dopamine neuron morphology and output are differentially controlled by mTORC1 and mTORC2"

**Supplemental Table 2. Raw values for HPLC measurements, related to Figures 8 and 9.**

| | **DA-Raptor WT** | | | | **DA-Raptor KO** | | | | **WT vs KO** |
|---|---|---|---|---|---|---|---|---|---|
| measurement | Mean | SEM | n (samples) | n (mice) | Mean | SEM | n (samples) | n (mice) | p-value/ test |
| Dorsal striatum **DA** (pmol/mm$^3$) | 101.1 | 3.225 | 6 | 3 | 30.83 | 2.951 | 10 | 5 | **<0.0001** Welch's t-test |
| Dorsal striatum **DOPAC** (pmol/mm$^3$) | 2.573 | 0.4943 | 6 | 3 | 0.4440 | 0.0664 | 10 | 5 | **0.0073** Welch's t-test |
| Ventral striatum **DA** (pmol/mm$^3$) | 47.77 | 3.149 | 6 | 3 | 17.32 | 2.736 | 10 | 5 | **<0.0001** Welch's t-test |
| Ventral striatum **DOPAC** (pmol/mm$^3$) | 1.880 | 0.4212 | 6 | 3 | 0.6213 | 0.1722 | 8 | 4 | **0.0292** Welch's t-test |
| | **DA-Rictor WT** | | | | **DA-Rictor KO** | | | | **WT vs KO** |
| measurement | Mean | SEM | n (samples) | n (mice) | Mean | SEM | n (samples) | n (mice) | p-value/ test |
| Dorsal striatum **DA** (pmol/mm$^3$) | 110.8 | 11.07 | 10 | 5 | 88.25 | 4.058 | 10 | 5 | 0.0809 Welch's t-test |
| Dorsal striatum **DOPAC** (pmol/mm$^3$) | 3.083 | 0.6679 | 10 | 5 | 2.170 | 0.3891 | 10 | 5 | 0.2566 Welch's t-test |
| Ventral striatum **DA** (pmol/mm$^3$) | 57.82 | 6.855 | 10 | 5 | 42.90 | 6.525 | 10 | 5 | 0.1323 Welch's t-test |
| Ventral striatum **DOPAC** (pmol/mm$^3$) | 2.541 | 0.4733 | 10 | 5 | 1.793 | 0.2639 | 8 | 4 | 0.2447 Welch's t-test |
